# MetaDIA: A Novel Database Reduction Strategy for DIA Human Gut Metaproteomics

**DOI:** 10.1101/2024.03.14.585104

**Authors:** Haonan Duan, Zhibin Ning, Zhongzhi Sun, Tiannan Guo, Yingying Sun, Daniel Figeys

## Abstract

**Background:** Microbiomes, especially within the gut, are complex and may comprise hundreds of species. The identification of peptides in metaproteomics presents a significant challenge, as it involves matching peptides to mass spectra within an enormous search space for complex and unknown samples. This poses difficulties for both the accuracy and the speed of identification. Specifically, analysis of data-independent acquisition (DIA) datasets has relied on libraries constructed from prior data-dependent acquisition (DDA) results. This approach requires running the samples in DDA mode to construct a library from the identified results, which can then be used for the DIA data. However, this method is resource-intensive, consumes samples, and limits identification to peptides previously identified by DDA. These limitations restrict the application of DIA in metaproteomics research.

**Results:** We introduced a novel strategy to reduce the search space by utilizing species abundance and functional abundance information from the microbiome to score each peptide and prioritize those most likely to be detected. Employing this strategy, we have developed and optimized a workflow called MetaDIA for analysis of microbiome DIA data, which operates independently of DDA assistance. Our method demonstrated strong consistency with the traditional DDA-based library approach at both protein and functional levels.

**Conclusion:** Our approach successfully created a smaller, yet sufficient database for DIA data search requirements in metaproteomics, showing high consistency with results from the conventional DDA-based library. We believe this method can facilitate the application of DIA in metaproteomics.

## Introduction

The microbiome encompasses a diverse array of microorganisms residing in different organisms, ecosystems, and environmental settings such as the human body, animals, plants, soil, water bodies, and various ecological niches[1, 2]. Metaproteomics serves as a tool for understanding the roles of proteins within these microbial communities[3]. Mass spectrometry-based proteomics aims to study all proteins in a sample. However, applying these techniques to the microbiome is challenged by its complexity. Without prior knowledge of the microbes present in a sample, metaproteomics relies on searching mass spectra against a large database, making the task of matching peptides and spectra notably challenging. Employing an iterative search strategy significantly reduces the search complexity in which the final search is against a database generated from previous searching results [4, 5]. The iterative strategy has been successfully used but only for the data acquired by data-dependent acquisition (DDA) mode[6, 7], Unfortunately, in DDA mode, only the most abundant precursor ions are selected for further inquiry, and lower abundant ones are overlooked[8].

In contrast, data-independent acquisition (DIA) uses a set of precursor isolation windows to collect all the fragments ions indiscriminately[9]. It has shown remarkable robustness, sensitivity, and reproducibility with fewer missing values[10]. DIA can be coupled with microLC enabling high-throughput analysis[11]. This makes it particularly suitable for conducting large-scale analyses. The DIA-PASEF[12] method integrates ion mobility separation with the DIA workflow, adding a fourth dimension of analyzing ion mobility to the traditional three-dimensional data set. This not only enriches the structural information of analytes but also enhances ion utilization efficiency leveraging the linear relation between ion mobility and mass-to-charge ratio. Another improvement in mass spectrometer scanning speed enables the utilization of smaller isolation windows in DIA, termed as narrow-window DIA[13]. This approach achieves comprehensive peptide precursor coverage and high quantitative precision and accuracy. In bioinformatics, the development of prediction software for peptide properties (theoretically predicted spectrum[14, 15], retention times[16–18]) enables the querying of DIA datasets without dependence on libraries generated by DDA. Those predicted libraries even showed better performance than the measured libraries[19]. Moreover, DIA-specific searching software such as DIA-NN[20, 21], MaxDIA[22], and Spectronaut have shown reliable results for the identification and quantification of peptides. The above advantages make DIA increasingly popular in proteomics. However, it is noteworthy that the benefits conferred by these techniques have not yet been fully extended to the field of metaproteomics. The main reason is that the inherent complexity of DIA data requires a much more constrained searching space compared with DDA data. To date, only a few metaproteomics studies have been done, and they were all compelled to use a spectral library derived from DDA data[23–25]. The DDA-derived method involves creating a spectral library from DDA runs for each sample, which is then used to interpret complex mass spectra from subsequent analyses. This approach requires multiple sample aliquots, extensive mass spectrometry resources and is limited to detecting peptides previously identified by DDA. Gladiator[26] uses DIA-Umpire[27] to assemble pseudo-DDA spectra from DIA data for microbiome samples. The method does not require a DDA-based spectral library for its operation, however, it still relies on spectrum-centric algorithms and does not fully exploit the potential advantages of DIA data.

Therefore, to leverage the benefits of DIA in metaproteomics, the searching space needs to be further reduced. In the previous DDA iterative strategy[7, 28], the high-abundant proteins (HAP) were used for the first search to infer the species that exist in the sample then all the proteins belonging to those species were then used for the subsequent search. However, this database remains overly extensive when compared to the number of identified peptides. Since the abundance of species within the microbiome shows significant disparity[29], the species identified should not be considered equally. The same applies to proteins and peptides. Proteins with high abundance and peptides with high detectability[30] or shared among various species are more likely to be detected. Here we report on a DIA workflow for metaproteomics, called MetaDIA, that relies on an annotated peptide database. This database comprises peptides that are anticipated to be detected, leveraging information on species abundance and protein abundance to score each peptide. We conducted a proof-of-concept experiment on human gut microbiome data generated by diaPASEF mode[23]. The peptide identification number and quantitative results obtained through our peptide library are comparable to those from the DDA-based library. Moreover, the species and functional information obtained from both methods are highly consistent.

## Materials and methods

### Reference peptide sequence with detectability score for human gut microbiome

The Unified Human Gastrointestinal Protein (UHGP) catalog, encompassing 4744 assembled genomes from the human gut microbiome, served as the reference database for this study[31]. Within this catalog, each protein sequence is uniquely associated with a distinct genome and is accompanied by detailed taxonomic and functional annotations. The detectability of peptides derived from these protein sequences was predicted using DeepDetect[30], a deep learning algorithm specifically designed for this purpose. This process involved in silico digestion of the protein sequences and subsequent assignment of a detectability score to each resultant peptide. Consequently, the peptide sequence reference database was enhanced by annotating each peptide with three key pieces of information: the genome identifier, the protein identifier, and the peptide’s detectability score. Please note that the database is structured on an identifier-centric organization. This means that peptides with identical sequences may be present within the database; however, as long as they are not from same genome and protein, they are distinguished by unique identifiers.

### Generation of FuncTax score

Firstly, identified peptides by MetaPep[32] are mapped to the UHGP database to establish peptide-genome associations. Subsequently, a greedy algorithm is employed to identify the minimal set of genomes that encompasses all peptide sequences, effectively reducing the complexity of the dataset. Following this, the intensity of each peptide is aggregated to infer genome abundance. The relative genome abundance will be used as the taxonomic score. To address the assignment of shared peptides, a razor strategy is adopted, analogous to the MaxQuant approach for protein inference[33]. Specifically, when a peptide is found in multiple genomes, it is attributed to the genome with the greater number of associated peptides. However, this typically results in approximately 1,000 genomes remaining, with many containing only a single peptide. The number substantially larger than that is found in a typical human gut microbiome which is around 200[29]. So, we only choose the most abundant species for subsequent analysis. The selection of species for consideration is further explored in the optimization section of the study.

For the functional score, we constructed a fixed table from the MetaPep project [32]. While building the database Metapep, the peptide identification was performed by the software MetaLab MAG[7], which provides quantifications of protein abundance. Those proteins are well annotated. Subsequently, the relative abundance of each Clusters of Orthologous Groups (COG) accession was computed. Samples comprising fewer than 1000 COG accessions were considered to be of low quality and consequently were omitted from the analysis. A total of 1,031 high-quality samples were retained for further evaluation. The mean of non-zero relative abundance of the COG accessions was then determined across these 1,031 samples, establishing a metric referred to the functional score.

The FuncTax score was obtained by multiplying two scores. In the case of peptides with the same sequence, their FuncTax scores were combined to give higher priority to shared peptides; the highest detectability score among them was utilized to ensure the inclusion of all possible peptides.

### Taxonomic and functional analysis

The taxonomic analysis is similar to the generation of genomic abundance score. The identified peptides are mapped to a database to establish peptide-genome associations. The database contains only the top 50 genomes. In our workflow, the peptide database was filtered out from the top 50 genomes. So, all the identified peptides were from the top genomes and thus can be used for the taxonomic analysis (the peptides added from MetaPep may not be used). In the DDA-based method, the peptide identified by the DDA library can be annotated to over 1,000 genomes even after using the greedy algorithm described above (method: generation of FuncTax score). However, we found that the top 50 genomes accounted for 79%-90% of peptides and 87%-92% of peptide intensity (Supplementary Figure 1). To simplify the comparison between the two methods, we discarded the small number of peptides that cannot be annotated to the top 50 genomes. Similarly, the razor strategy is used to process peptides shared by multiple genomes. Finally, the intensity of each peptide is aggregated to infer genome abundance.

For functional analysis, a protein abundance was firstly generated using the same strategy as taxonomic analysis. The proteins in the UHGP database have been extensively annotated thus the protein abundance can be further interpreted into functional abundance.

### Deepdetect software configuration

Protein digestion was simulated using Trypsin with the following parameters: a maximum of two missed cleavages, and peptide lengths ranging from 7 to 50 amino acids. Default settings were applied for all other parameters.

### DIA software configuration

DIA-NN (version 1.8.1) was used to process all the DIA data in this study. Maximum mass accuracy tolerances were set to 10 ppm for both MS1 and MS2 spectra. The --relaxed-prot-inf option was used for library-free searching. The --no-maxlfq option was used to disable the normalization for the quantification benchmark experiment. All other settings were left default. The precursor matrix containing the peptide information was used for taxonomic and functional analysis.

### Metaproteomic datasets

The dataset used for optimizing workflow is sourced from a published study and shared by the authors[23]. The dataset for evaluating accuracy is from in-house samples. *Blautia hydrogenotrophica* (DSM 101114; Leibniz Institute DSMZ- German collection of microorganisms and cell cultures) was cultured in LB broth. The human stool was collected from a healthy adult volunteer at the University of Ottawa, Ottawa, ON, CAN. The protocol (# 20160585-01H) was approved by Ottawa Health Science Network Research Ethics. The protein extraction and digestion were performed as described previously[34]. Peptide concentrations were measured using Thermo Scientific Pierce Quantitative Colorimetric Peptide Assays according to the manufacturer’s directions.

The in-house samples were then analysed using an UltiMate 3000 RSLCnano system (Thermo Fisher Scientific, USA) coupled to an Orbitrap Exploris 480 mass spectrometer (Thermo Fisher Scientific, USA). Peptides were loaded onto a tip column (75 μm inner diameter ×15 cm) packed with reverse phase beads (3 μm/120 Å ReproSil-Pur C18 resin, Dr. Maisch HPLC GmbH). A 60 min gradient of 5 to 35% (v/v) from buffer A (0.1% (v/v) formic acid) to B (0.1% (v/v) formic acid with 80% (v/v) acetonitrile) at a flow rate of 300 μL/min was used. The mass spectrometer was in data-independent mode covering the mass range of 380–980 m/z with 10 m/z isolation windows.

### Availability of the pipeline

The whole pipeline is available for use at https://github.com/northomics/MetaDIA

## Result

### MetaDIA Workflow overview: Taxonomy- and function-guided construction of peptide database for metaproteomics

Here we propose a new workflow for DIA based metaproteomics called MetaDIA. MetaDIA is a multistep workflow that systematically reduces the search space for DIA searching. At its basis, it relies on a combination of taxonomic abundance, functional abundance as a proxy of protein levels, and peptide detectability ultimately enabling DIA searching without the need for DDA results. Briefly, in the first step, we created a new database of peptides, called MetaPepDetec, obtained by in silico digestion and detectability prediction of the Unified Human Gastrointestinal Protein (UHGP, 4744 genomes) database into peptides[30, 31]. Then each peptide is annotated with a FuncTax score (Figure 1). Both the FuncTax and the detectability scores are used to reduce the peptide database.

**Figure 1.**
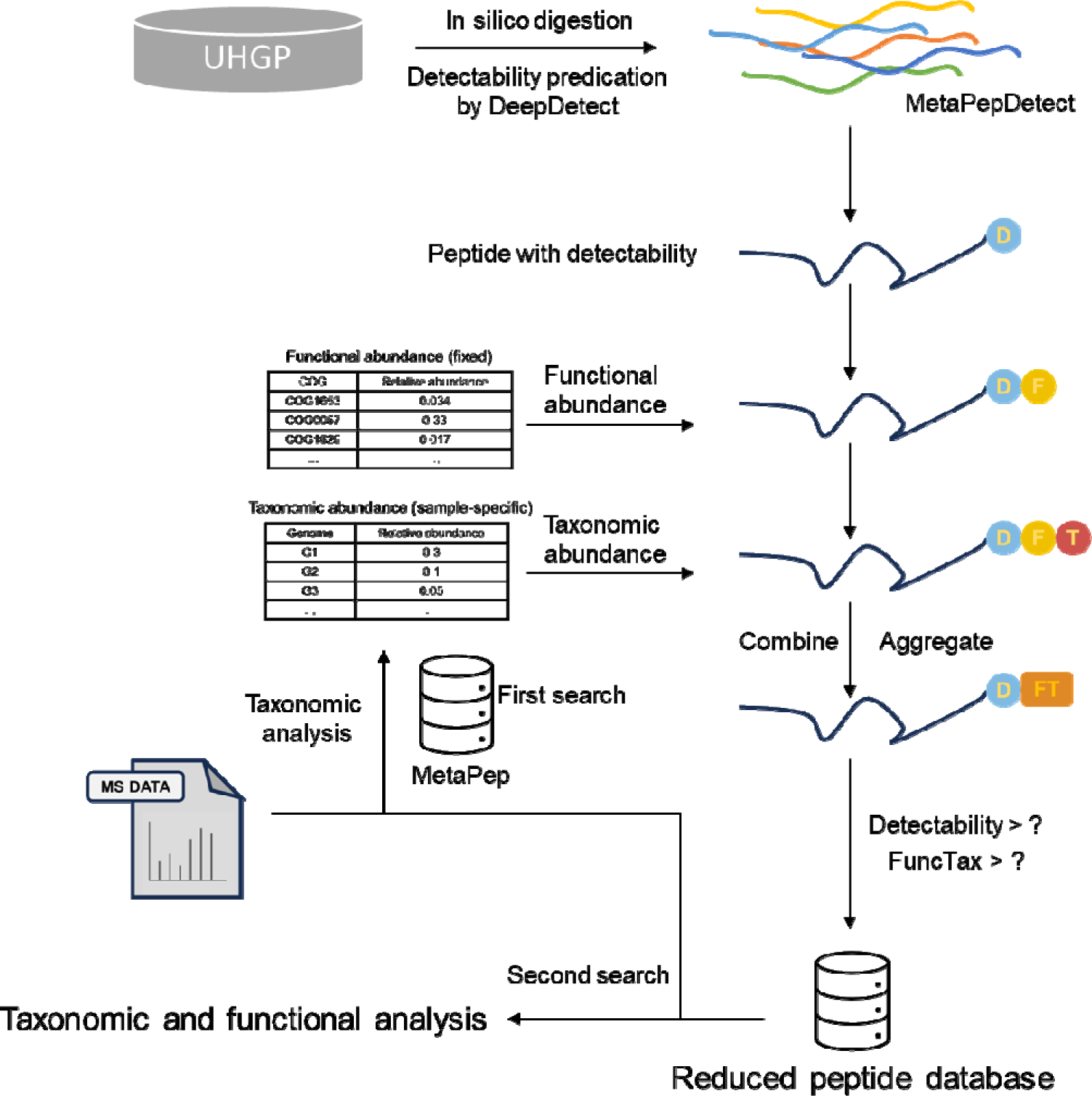
The flowchart for the MetaDIA. All proteins in Unified Human Gastrointestinal Protein (UHGP) database were firstly in silico digested into peptides. The detectability of each peptide was predicated by DeepDetect algorithm. Following this prediction, each peptide was assigned a functional score and a taxonomic score, derived from a predetermined functional relative abundance table and a sample-specific taxonomic relative abundance table, respectively (method). The FuncTax score was calculated by multiplying the two scores. For peptides with identical sequence, their FuncTax scores were aggregated to prioritize shared peptides; the maximum of their detectability scores was used to ensure the inclusion of all potential peptides. The detectability and FuncTac scores are both used for filtering peptides. The reduced peptide database was used for a second search.

The FuncTax scores for each peptide in the MetaPepDetec are calculated using information from the MetaPep database [32]. MetaPep is a core peptide database compiling peptides previously identified in the published human gut metaproteomics studies. The information from MetaPep was used to create a static table of COG relative functional abundances and a sample-specific table of taxonomic relative abundances (method). We noted that despite significant differences in the species composition of gut bacteria among different individuals, their functions are remarkably similar[35]. Therefore, functional abundance hierarchy information could act to estimate the likelihood of a protein being observed. We analyzed the search results used to construct the MetaPep database which contains 2,134 raw files and 415 individuals[32] (Method). The functional ranking among various samples exhibits a strong correlation (Supplementary Figure 2a and 2b). We observed a stable pattern in the functional hierarchy of human gut bacteria: abundant functions consistently remain high, while scarce functions persistently stay low across all samples (Supplementary Figure 2c and Supplementary File 1). The sample-specific table of taxonomic relative abundances was generated by searching the DIA data against MetaPep[32]. The identified peptides and their quantitation were used to create the table (Method). The FuncTax score for each peptide is calculated by multiplying the taxonomic score for its taxonomic annotation and the functional score of its functional annotation. For peptides with identical sequences, their FuncTax scores were aggregated thereby leading to shared peptides having a higher ranking.

In the last step, sample-specific reduced peptide database is generated by filtering MetaPepDetec using the FuncTax score and the detectability score (Figure 1). The final search of the DIA data is done against the reduced peptide database. To validate the efficiency of our peptide ranking method, peptides were sorted by FuncTax score and partitioned into equal-sized subsets based on their percentile rank (e.g., top 0-5%, 5-10%, …, 35-40%). Each subset was subjected to database searching with uniform parameters. We observed a decline in the number of peptides identified as the percentile ranking of the subsets decreased (Figure 2). The decreasing trend suggested our ranking method effectively prioritizes peptides with a higher probability of detection.

**Figure 2.**
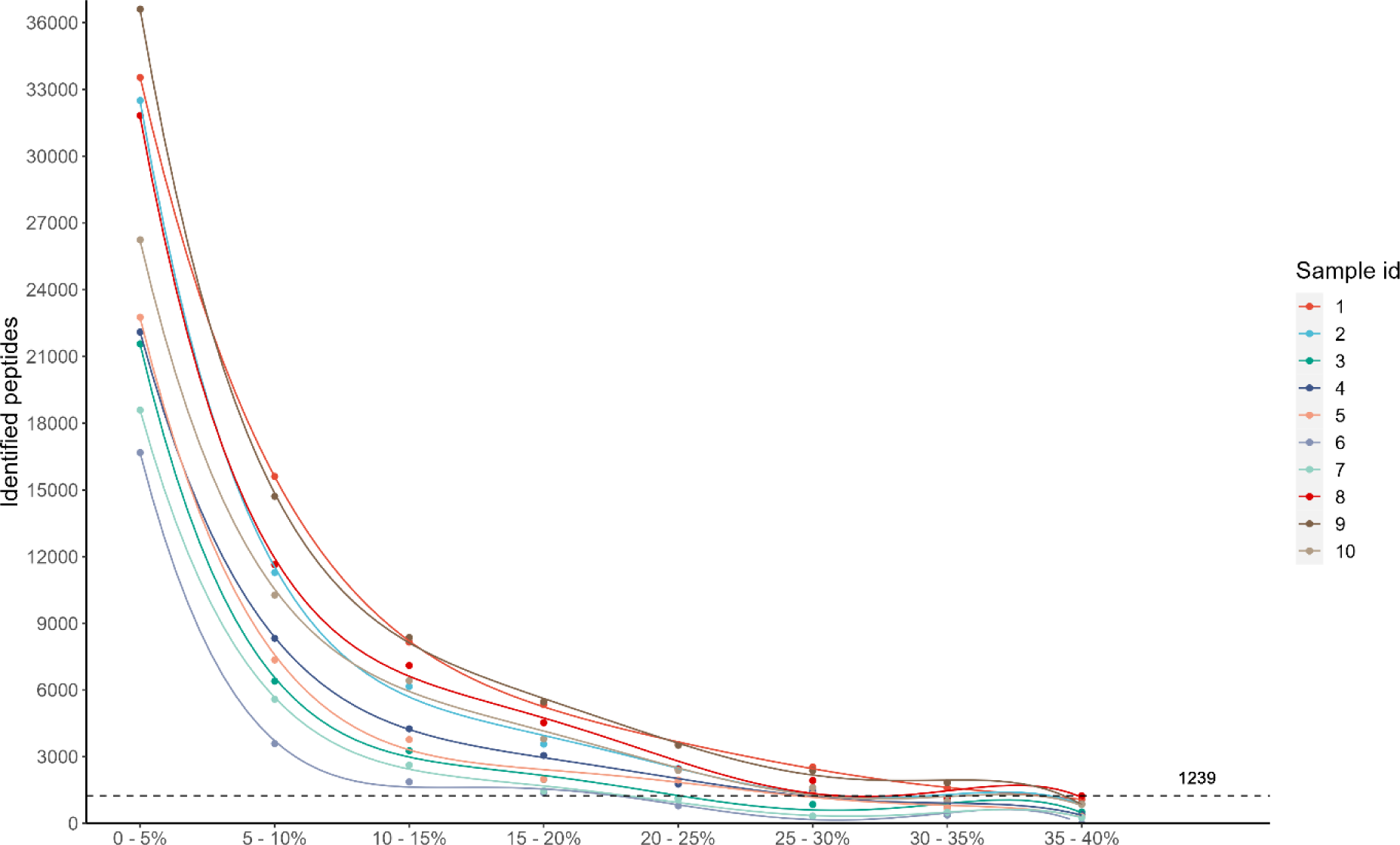
The number of peptides identified from each subset. Ten samples were tested in the experiment. For constructing the peptide database, the top 100 genomes were considered; the detectability threshold was set at 40%. Each subset contains around 400,000 peptides. Peptide identification was performed by DIA-NN under same conditions. The maximum identification from the last subset was heighted in the figure.

### Optimized MetaDIA parameters reduces the database size

We explored whether the number of microbes in the reduced peptide database, the threshold for FuncTax score and the threshold for detectability score influenced the identification of peptides. We explored the impacts of the parameters using the DIA data from 10 different human gut microbiome samples previously reported[23] (Supplementary File 2, sample information).

In particular, we first tested effect of the number of microbes (genomes) ranging from 50 to 150 and FuncTax score ranging from top 1% to 40% (Figure 3). We keep the detectability threshold at the top 40% in this experiment which is suggested by the author of Deepdetect[30]. Interestingly, no mater how many genomes we choose, the size of the reduced peptide database had the strongest effect on the number of identified peptides. The identification number plateaued once the reduced peptide database size reached around 1.6 million entries (Figure 3a and 3b, Supplementary Figure 3 and 4), corresponding to a FuncTax score threshold of 40% for 50 genomes, 20% for 100 genomes and 15% for 150 genomes respectively. We compared the three different reduced peptide databases, which led to consistent peptide identification results (Figure 2c and 2d, Supplementary Figure 5 and 6). In our previous studies, we observed that low-abundance species were underrepresented[8]. In this context, we chose to focus on the top 50 genomes to prioritize high-abundance genomes. It is important to note that this cut-off is a variable parameter that can be adjusted according to the specific objectives of different studies.

**Figure 3.**
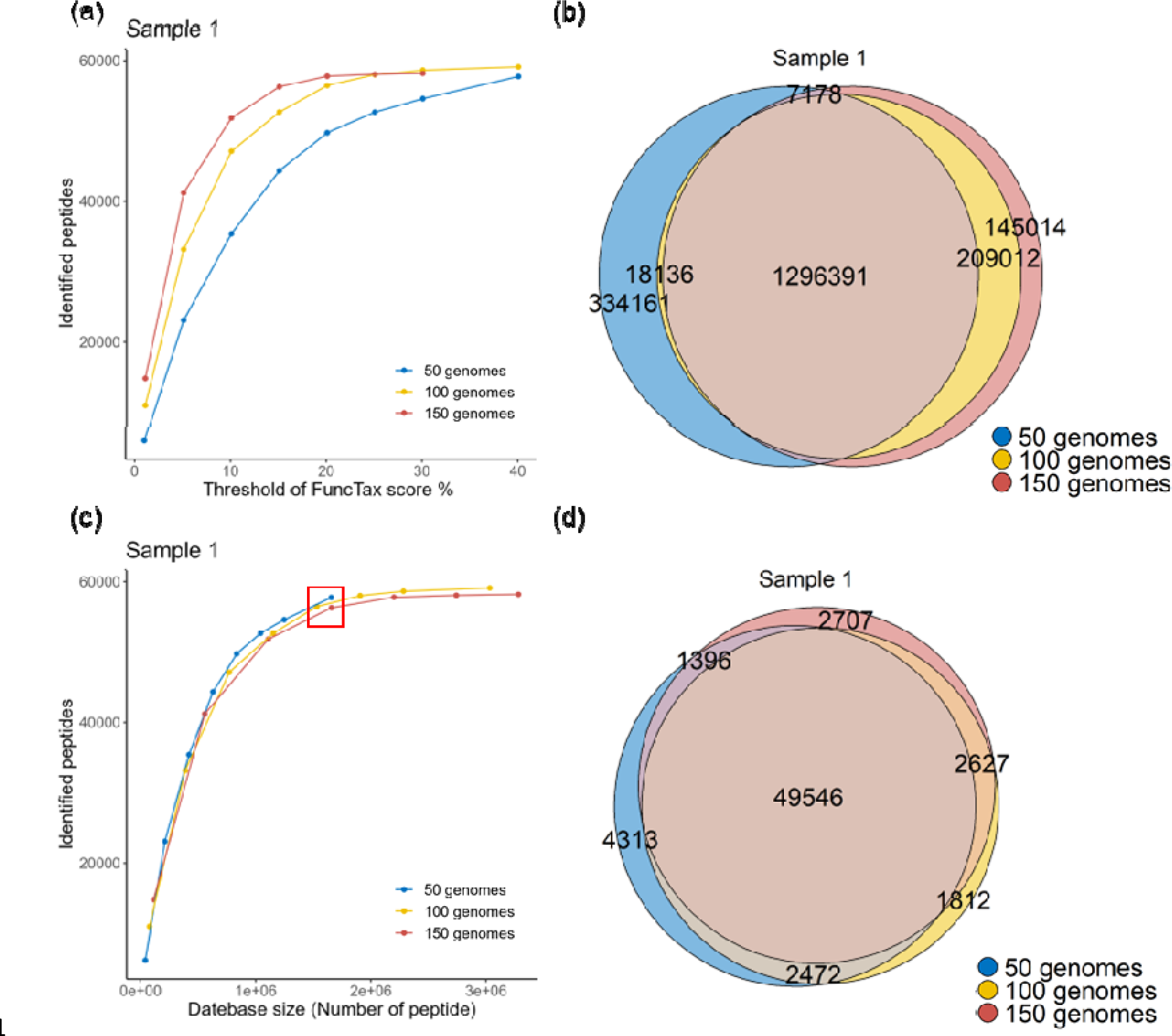
Optimization for genome number and FuncTax score (Sample 1 was shown. For the other samples, please see Supplementary figures). Peptides from n (50, 100, 150) genomes were ranked by the FuncTax score and top x% (1-40 for 50 and 100 genomes; 1–35 for 150 genomes) peptides was used as database. (a) Number of identified peptides against database percent. (b) Number of identified peptides against database size. The inflection point has been highlighted with a red box. (c) The overlap of the reduced peptide database and (d) identified peptide when taking top 40% peptides for 50 genomes, top 20% for 100 genomes and top 15% for 150 genomes as database. Peptide identification was performed by DIA-NN under same conditions.

Subsequently, we explored whether the detectability threshold impacted the number of peptides identified. While the recommended threshold by the author of Deepdetect is 40%, we explored thresholds ranging from 40% to 10%. We observed that a threshold of 25% was the point at which the number of identifications began to decrease significantly (Figure 4a). However, both the database size and the search time decreased substantially (Figure 4b). Comparing the identification results at thresholds of 25% and 40%, we found a substantial overlap (Figure 4c).

**Figure 4.**
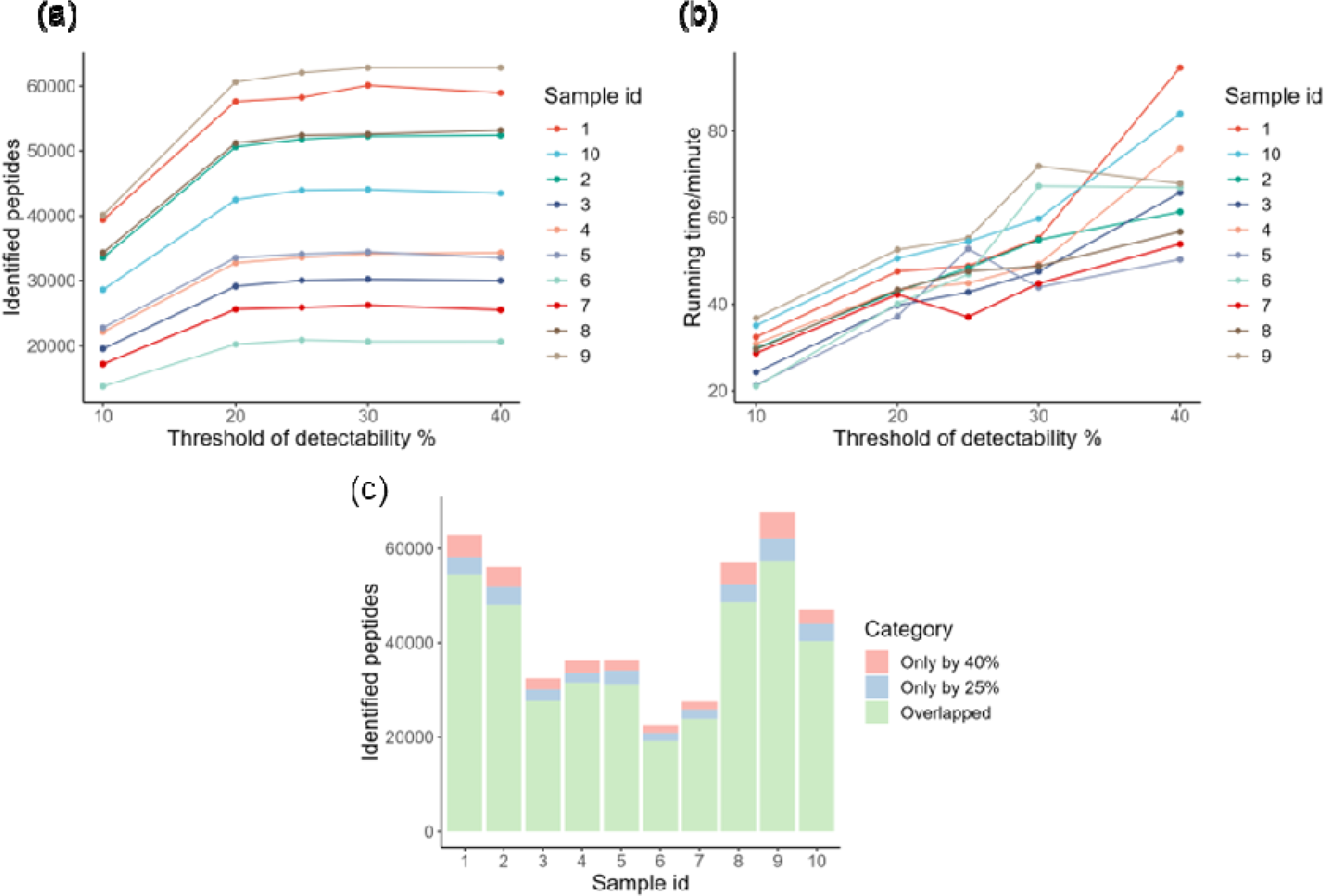
Optimization for detectability threshold. (a) The number of peptides identified and (b) the searching time under detectability threshold from 10% to 40%. (c) The overlap of peptides identified by top 25% and top 40% of the database. Peptide identification was performed by DIA-NN under same conditions.

Therefore, we selected a 25% threshold for detectability. Based on this analysis, we proceeded with peptides ranking in the top 40% by FuncTax score and the top 25% by detectability. Given that these two scores are entirely uncorrelated, applying both filters effectively reduced the database to one-tenth of its original size (25% times 40%, Supplementary Figure 7). After applying these optimized parameters, approximately 1 million peptide sequences remain in the reduced database.

### MetaDIA maintains accuracy in DIA peptide identification

We next explored whether the enrichment of high abundant and highly detectable peptides in our reduced database impacted the accuracy of peptide identification when applying the false discovery rate strategy. To evaluate this, we conducted a benchmark experiment using three samples: a human gut microbiome sample (A), a *Blautia hydrogenotrophica* sample (C), and a 50:50 mixed sample of the two (B) (Figure 5a). *Blautia hydrogenotrophica* was selected due to its absence in the microbiome sample used here and its minimal peptide overlap with the microbiome sample. Each sample was subjected to triplicate DIA measurements. Sample A and B were analyzed using the reduced peptide database generated by our workflow with optimized parameters of top 40% FuncTax score and top 25% detectability score, whereas sample B and C were searched against species-specific protein databases derived from NCBI (Genome assembly ASM15797v1). In the first search against the peptide database, 32,624 unique peptides were identified. Of these, 1,952 peptides also present in the *Blautia hydrogenotrophica* database were excluded. Further, peptides unique to each sample were removed, leaving 27,830 peptides identified in both sample A and B. Ideally, the peptide abundance ratio between samples A and B should approximate 2. In the second search, 14,821 unique peptides were identified. Among these, 10,995 peptides were unique to the *Blautia hydrogenotrophica* database and were found in both samples B and C. The expected ratio between samples B and C should be around 0.5. We found whether using a protein database or a peptide database, the ratios of peptides identified in both searches closely aligned with the expected values (Figure 5b). This suggests that the employment of our reduced peptide database does not significantly affect the accuracy of peptide identification, thereby supporting its use in peptide identification workflows with a controlled FDR.

**Figure 5.**
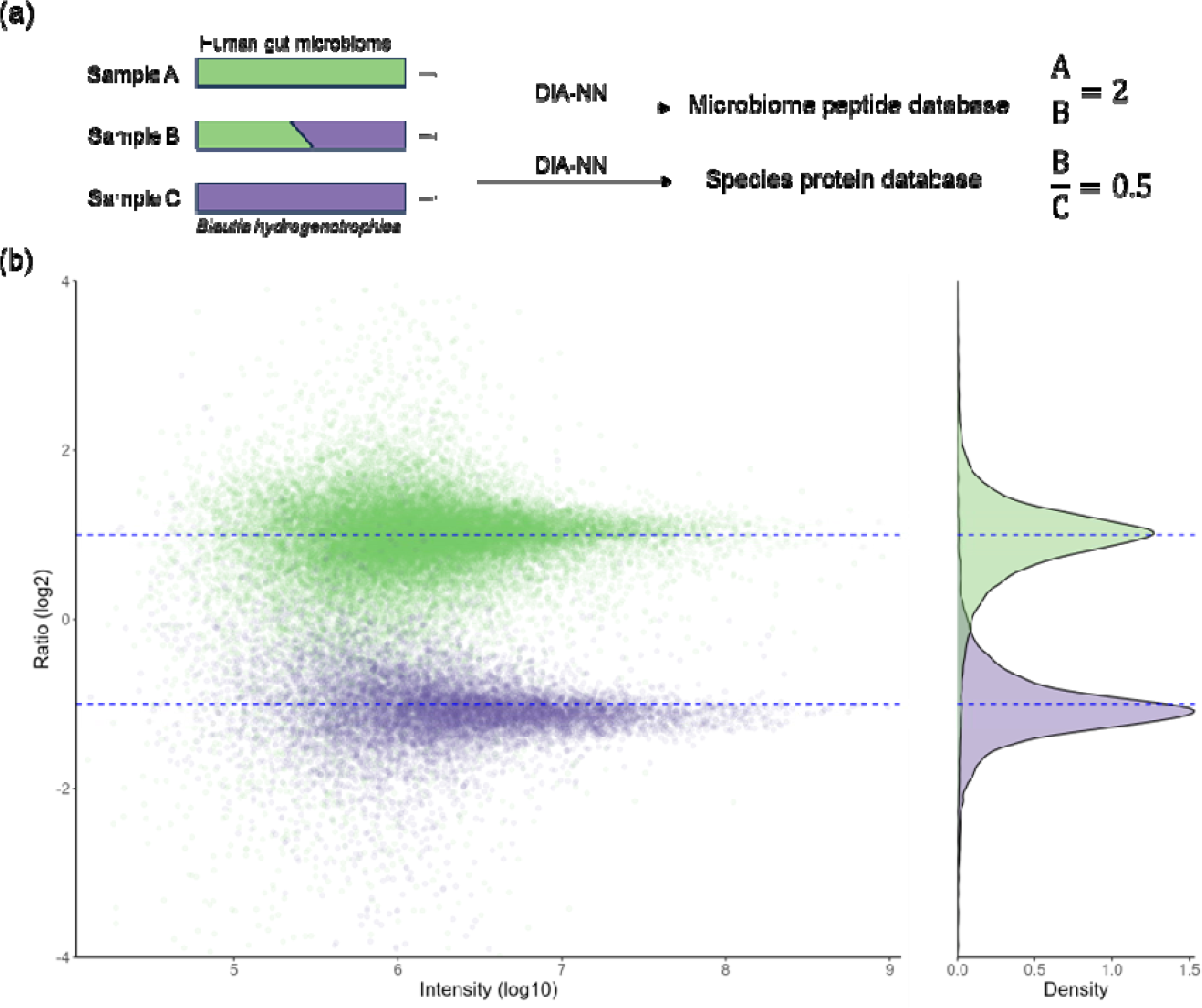
Benchmark experiment for peptide identification. (a) The experimental design. Each sample was subjected to triple-run measurements (b) Log-transformed ratios are plotted as a function of peptide intensity for n = 27,830 (green) microbiome peptides and n = 10,995 (purple) *Blautia hydrogenotrophica* peptides. The point density for ratio was plotted at right. Dashed lines indicate the expected ratio. Peptide identification was performed by DIA-NN under same conditions. The intensities derived from various charge states of the same peptide were aggregated.

### MetaDIA yields consistent peptide and protein identification results with DDA based strategies

We next evaluated whether MetaDIA performed similarly to a conventional DDA-based library for DIA data analysis. The DIA data and corresponding DDA-based library were obtained from a published study[23]. We found that the MetaDIA provided identification numbers comparable to those obtained through the DDA-based library (Figure 6a). Notably, in certain instances, such as with samples 8 and 9, the MetaDIA surpassed DDA library in the number of identifications. The initial step in our workflow involves searching the raw data against MetaPep, which leverages the results from an open search strategy, thereby encompassing modified peptides not included in the original database. Subsequent integration of peptides identified by MetaPep into a refined peptide database resulted in a marked increase in identification rates (Figure 6a).

**Figure 6.**
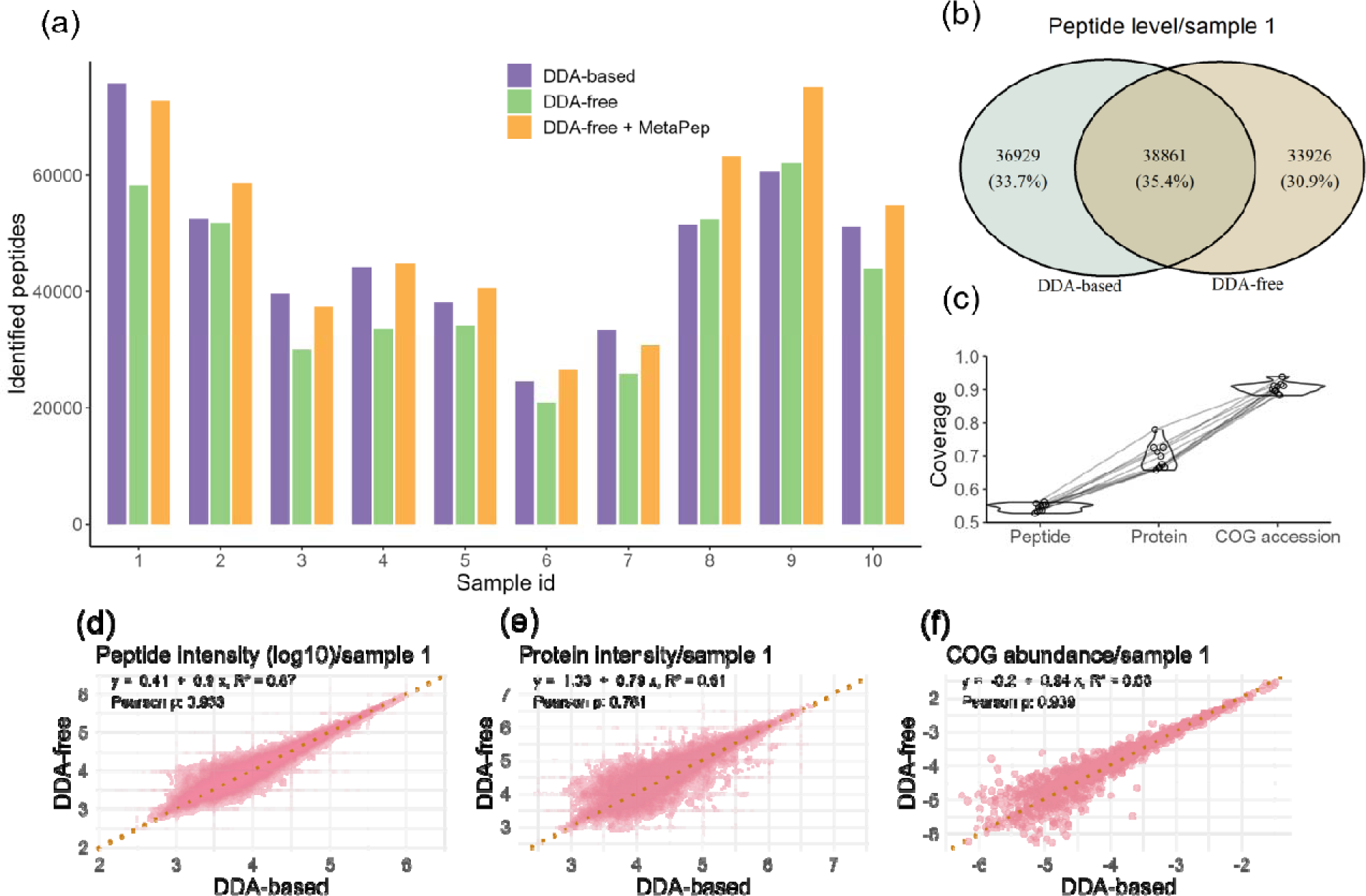
Comparison between the DDA-based method and DDA-free method. (a) The peptide identified by each method. (b) The overlap of peptide identified in sample 1 by each method. (c) Coverage of peptides, proteins and cog accessions identified by DDA-based method with those found using DDA-free method. The intensity correlation of the overlapped peptides (d), proteins (e) and COG accessions in sample 1. The dashed line indicates y = x. For DDA-based method, the peptides identified as derived from human proteins are removed.

Over 50% of peptides identified from the DDA-based library were also identified by MetaDIA (Figure 6b and Supplementary Figure 8). The divergence in unique identifications between the two methods may be attributed to inherent differences between DDA acquisition and DIA acquisition. Upon examining the quantification results of those peptides found by both methods, we observed a significant consistency in the outcomes, with a Pearson coefficient above 0.9 (Figure 6d and Supplementary Figure 9). It is worth noting that the fragment ions used for quantification in the DDA-based library correspond to actual DDA acquisitions. In contrast, MetaDIA uses theoretical spectra that are predicted from peptide sequences. The high degree of agreement between the quantification results underscores the reliability of the MS-Simulator algorithm which is employed by DIA-NN for spectra prediction [14].

At the protein level, our findings revealed greater consistency in identification compared to the peptide level (Figure 6c). Around 70% of proteins found by the DDA-based library can be found by MetaDIA. The overlap on protein level reinforces the reliability of the identifications and indicates that a significant subset of proteins is consistently identified by both methods despite differences at the peptide level (Supplementary Figure 10). Proteins like GYG000002545_00035 had greater sequence coverage and higher detection intensity with the DDA library, while others like MGYG000002272_00452 showed higher coverage and intensity with MetaDIA. Given that the quantification of a protein is derived from different subsets of peptides in these two methods, we observed reduced consistency of quantification in the protein level between the methods, as reflected by Pearson correlation coefficients of approximately 0.7 (Figure 6e and Supplementary Figure 11). However, it is important to note that in most proteomic studies, the primary interest lies in the differential abundance of the same protein across various samples. Therefore, it is crucial that we use the same fragment ions to quantify a protein. In this regard, the inconsistencies in protein quantification between the two methods do not undermine the utility of either approach. The substantial overlap in peptide and protein identification by both methods suggests a robust cross-validation of both methods. Then we annotated the proteins using COG accessions and calculated their relative abundances. Our analysis revealed that approximately 90% of the COG accessions identified by the DDA-based method were also covered by our MetaDIA (Figure 6c). Furthermore, the Pearson correlation coefficient for the relative abundance of COG accessions exceeded 0.9, with a stronger correlation for those COG accessions that were highly abundant (Figure 6f and Supplementary Figure 12).

### MetaDIA provides taxonomic profiles highly similar to those obtained from searching DDA-libraries

We verified whether both methods had a high degree of similarity in the taxonomic composition. We did comparative analysis of microbiome composition across different taxonomic levels using the result from both methods. Our findings indicate that there is a significant linear correlation between the compositions identified by both methods, with the degree of correlation strengthening at higher taxonomic levels (Figure 7a and b, Supplementary Figure 13). The two methods showed remarkably consistent taxonomic composition at the genus level with a Pearson coefficient above 0.98 across all the samples tested. Even Sample 9, which displayed the lowest correlation, demonstrated a substantial degree of consistency between the two methods. To underscore the consistency, we have provided a detailed visualization of the taxonomic composition for Sample 9 (Figure 7c and d, Supplementary Figure 14) The species compositions observed by MetaDIA in these ten samples differed significantly as expected, indicating that our database and taxonomic analysis have the capability to identify a diverse range of microbiota (Supplementary Figure 14 and Supplementary File 3: searching result). The most abundant species identified in the ten samples have been previously reported as high-abundance species in the human gut microbiome[36–40]. Except for *Phocaeicola dorei* which were identified as the top species in sample 2, 5 and 10, the other top species were all unique to each sample.

**Figure 7.**
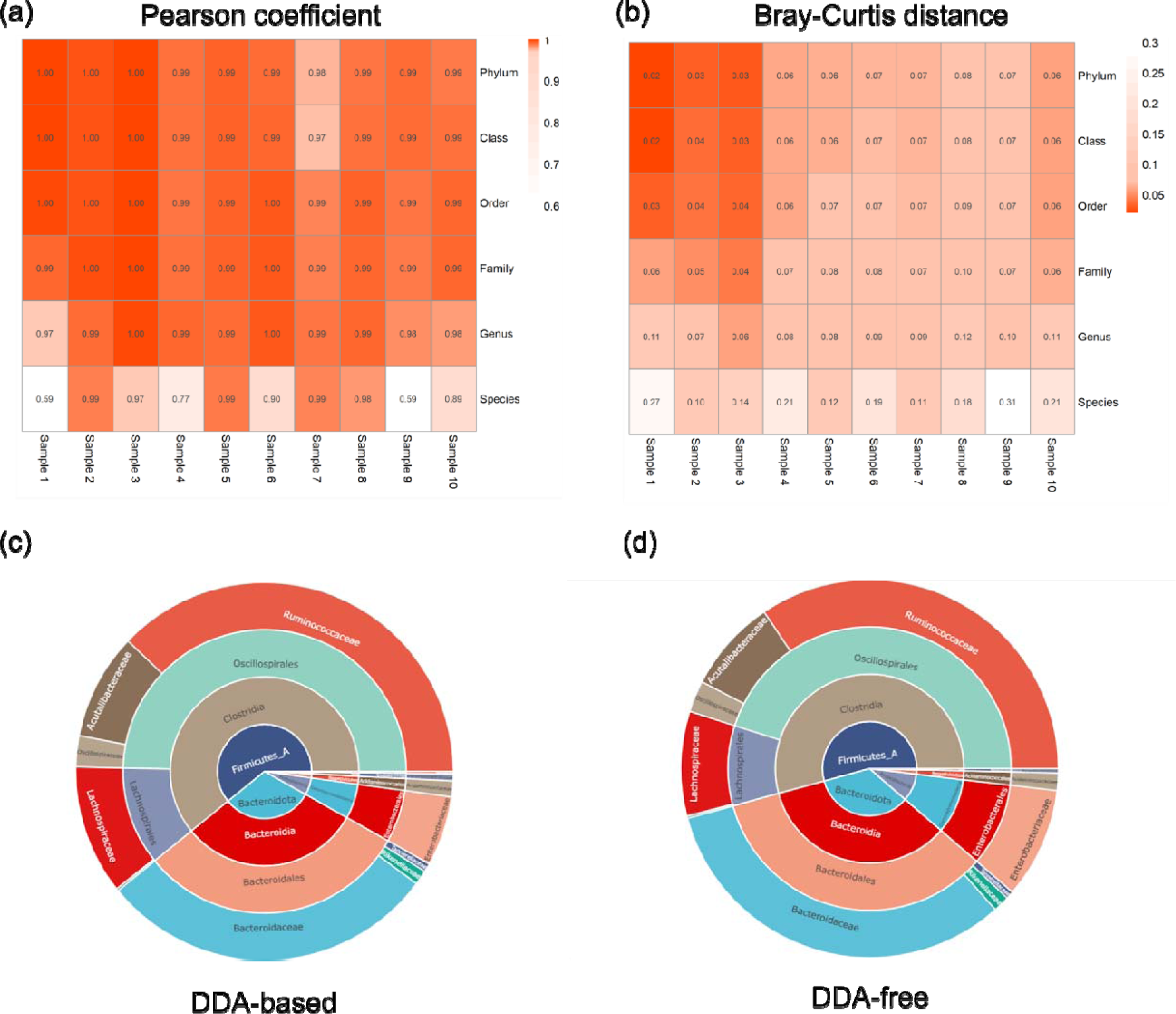
Comparison of the taxonomic composition between the DDA-based method and DDA-free method. Pearson correlation (a) and Bray-Curtis distance (b) analysis between DDA-based method and DDA-free method on different taxonomic levels from Phylum to Species. The relative taxonomic abundance was used for the analysis. In the correlation analysis, taxonomic categories that were unique to one method were imputed with a value of zero. The taxonomic composition (Phylum to Family) of sample 9 derived from DDA-based method (c) and DDA-free method (d).

### MetaDIA is universally applicable to different types of DIA, including DIA-PASEF

To further validate the versatility and applicability of our proposed metaproteomic workflow, we extended our analysis to a diverse set of 79 DIA datasets obtained from a published study[23]. This dataset encompasses samples from 62 individuals, featuring replicate injections, quality control (QC) samples, and pooled samples (Supplementary File 2: Sample information). We applied MetaDIA to this extensive dataset and compared the results with the conventional DDA-based approach. Remarkably, the number of peptides identified by both methods demonstrated a close equivalence, reinforcing the robustness and universal applicability of our metaproteomic workflow (Figure 8). Validating our method across diverse samples enhances confidence in its effectiveness and consistency, demonstrating its potential for widespread adoption in metaproteomics research.

**Figure 8.**
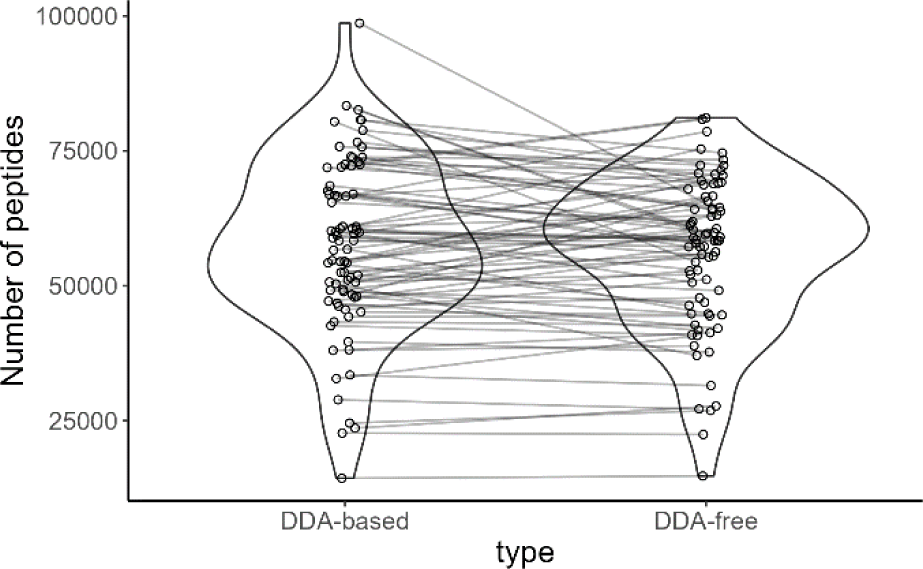
Number of peptides identified by both methods with mean value from 79 diaPASEF samples. For DDA-based method, the peptides identified as derived from human proteins are removed.

## Discussion

We propose a novel workflow for DIA data analysis from human gut microbiome called MetaDIA. The approach aims at prioritizing peptides with a higher likelihood of detection based on their detectability, taxonomic and functional scores.

MetaDIA is entirely devoid of DDA, thereby circumventing the drawbacks of DDA-based methods. Not only does this approach save time and resources, but it also enables the creation of a tailored database for each sample. In contrast, DDA-based methods typically rely on a single pooled sample to generate a library. For instance, Gomez *et al*.[25] used a pooled sample to represent 12 individual mice, while Sun *et al*.[23] did so for a cohort of 62 individuals. However, such a pooled sample may not effectively represent every sample. In our study, the ten samples showed highly diverse taxonomic composition (Supplementary Figure 13). To increase the sampling depth for the pooled sample, Sun *et al.*[23] had to fractionate the pooled sample into 30 portions and Gomez *et al*.[25] repeatedly injected the pooled sample 10 times. Moreover, utilizing a static library to search various samples may potentially compromise the accuracy of peptide identification, as it includes peptides from the pooled samples that are absent in the specific sample under investigation.

In MetaDIA, we pre-defined the range of genomes for each microbiome sample (50 genomes in this study). This approach not only enabled us to narrow the search space but also to mitigate the issues associated with protein inference that arise from common peptides. When assigning peptides to proteins, we confined our consideration to the genomes within the predefined range rather than the entire dataset. This strategy significantly reduced the incidence of common peptides.

Although MetaDIA is currently focused on the human gut microbiome, we foresee that it can be extended to other types of microbiomes, such as those in animal intestines, environmental microbiomes when using an appropriate bait database. A database similar to MetaPep could be constructed for other microbiomes.

## Conclusion

In conclusion, we introduced a new strategy to prioritize peptides with a high probability of detection. This strategy simulates protein digestion procedures in silico and uses taxonomic and functional information to infer the peptide abundance. MetaDIA is a fully DDA-free workflow and provides a user interface to change the different parameters. We compared the performance of MetaDIA with the DDA-based library and observed a high degree of consistency. We further validated our method across a DIA-PASEF dataset with 79 samples, thereby confirming its wide applicability. We believe that our approach will help the application of DIA in metaproteomics.

## Author information

### Authors and Affiliations

School of Pharmaceutical Sciences, Faculty of Medicine, University of Ottawa, Ottawa, ON K1H 8M5, Canada

Haonan Duan, Zhibin Ning, Zhongzhi Sun & Daniel Figeys

Department of Biochemistry, Microbiology and Immunology, Faculty of Medicine, University of Ottawa, Ottawa, ON K1H 8M5, Canada

Haonan Duan & Zhongzhi Sun

Westlake Center for Intelligent Proteomics, Westlake Laboratory of Life Sciences and Biomedicine, Hangzhou, Zhejiang Province, 310030, China

Tiannan Guo & Yingying Sun

School of Medicine, School of Life Sciences, Westlake University, Hangzhou, Zhejiang Province, 310030, China

Tiannan Guo & Yingying Sun

Research Center for Industries of the Future, Westlake University, 600 Dunyu Road, Hangzhou, Zhejiang, 310030, China

Tiannan Guo & Yingying Sun

### Contributions

HD, ZN, ZS, TG, YS and DF designed the study. HD performed the experiments. HD, ZN, DF analyzed the data. TG and YS shared the raw data. HD and DF wrote the manuscript. All authors read and approved the final manuscript.

### Corresponding authors

Correspondence to Daniel Figeys

## Acknowledgements

The authors acknowledge the assistance of OpenAI’s ChatGPT in grammar correction and clarification of this article.

## Ethics declarations

### Consent for publication

Not applicable

---

**Supplementary Figure 1.**
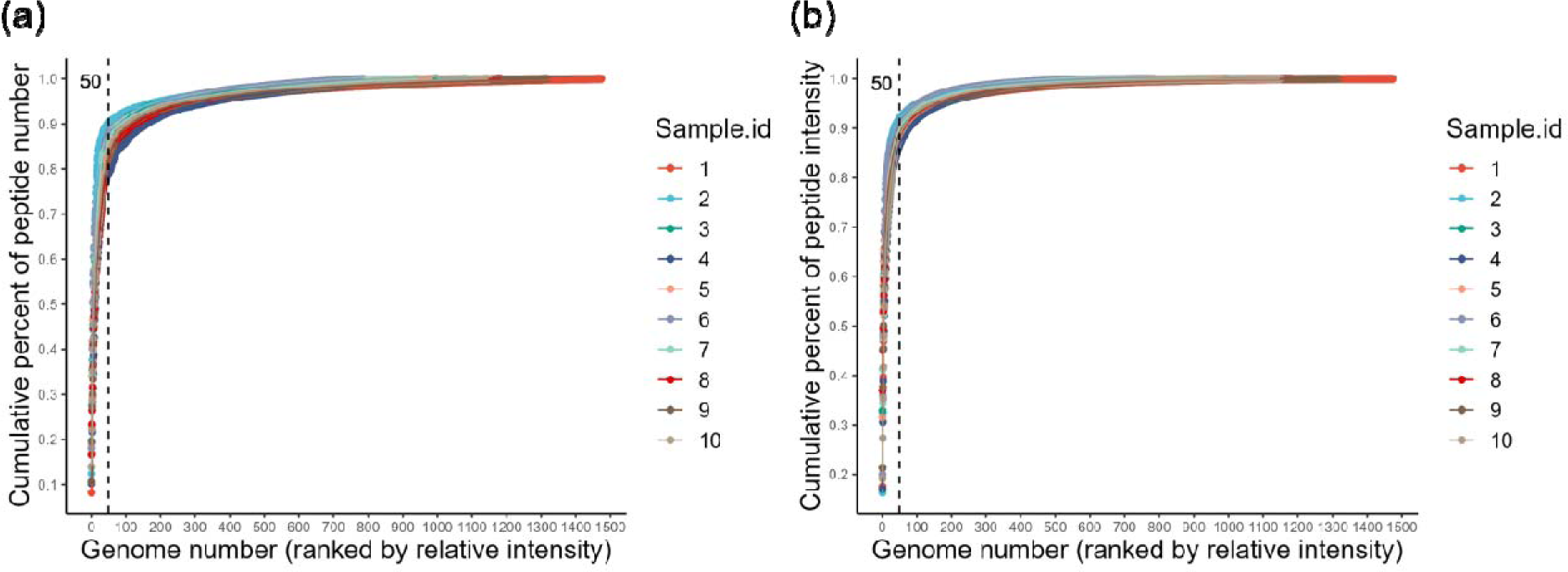
Cumulative contribution of identified peptides (a) and intensity (b). Figure were plotted against the number of genomes. Genomes are ordered by decreasing peptide count.

**Supplementary Figure 2.**
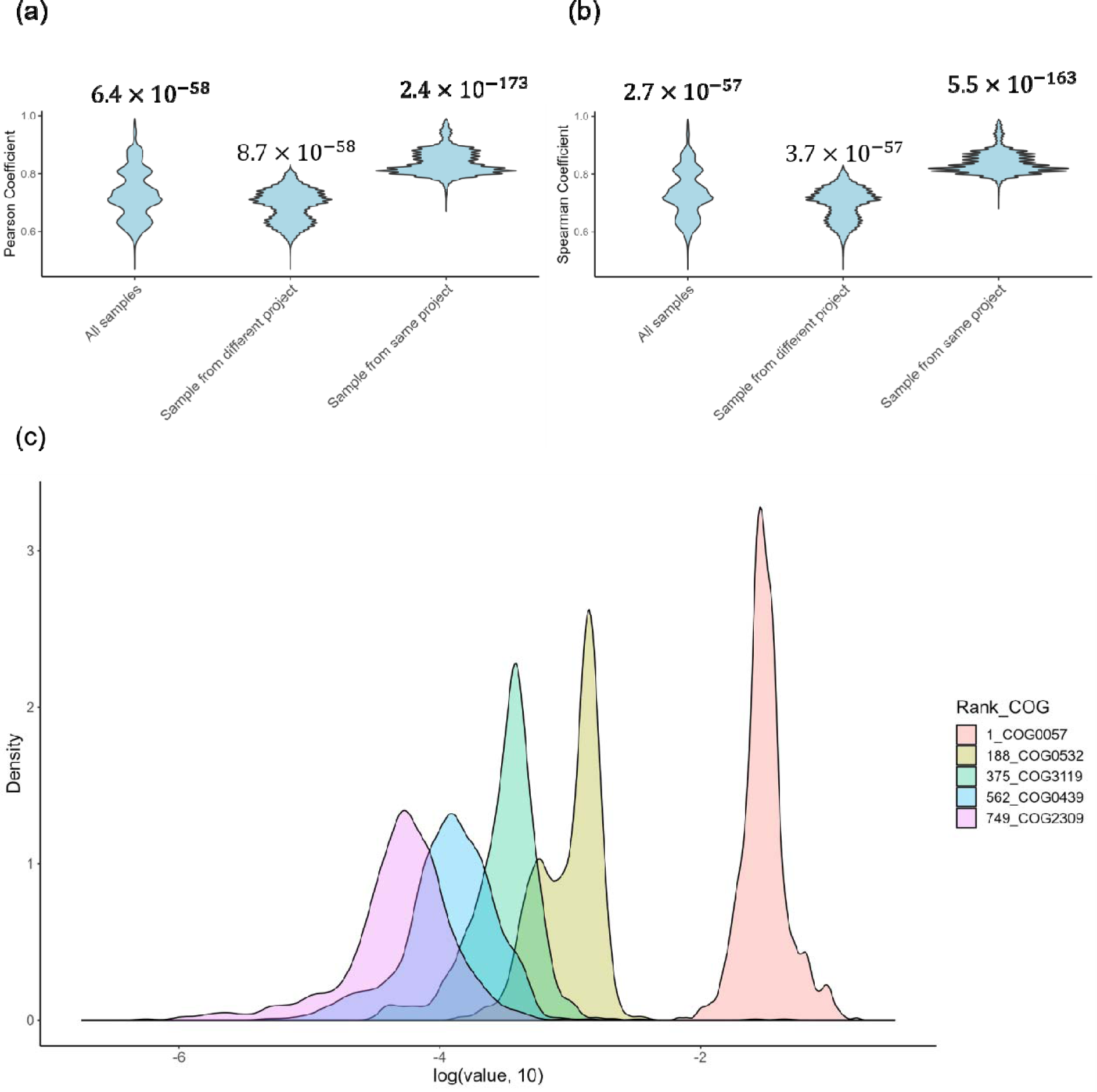
The functional correlation across samples in MetaPep. The Pearson coefficient (a) and Spearman coefficient (b) on COG accessions. Sample pairs from same project are likely from the same individuals and plotted in different groups. The average p value are plotted above each groups. (c) The distribution of relative abundance of COG accession across samples in MetaPep. The legends shows the rank and name of COG accessions. COG accessions present in more than 95% of the samples were retained. A total of 750 COG accessions remained and were subsequently sorted based on their functional abundance scores. Five COG accessions were selected for density plot at evenly spaced intervals from this ordered list.

**Supplementary Figure 3.**
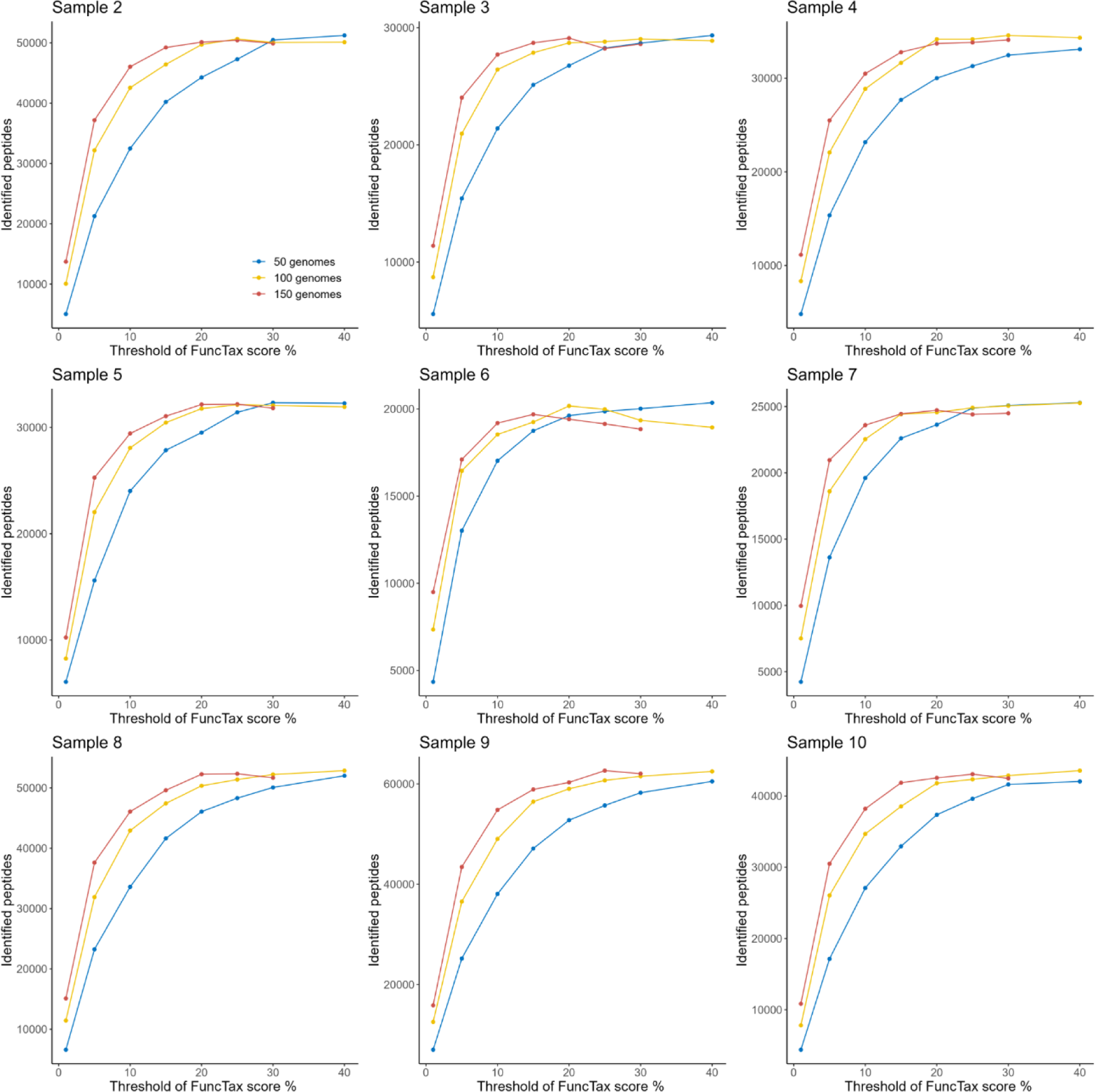
Optimization for genome number and FuncTax score (Sample 2-10). Number of identified peptides against database percent. Peptides from n (50, 100, 150) genomes were ranked by the FuncTax score and top x% (1-40 for 50 and 100 genomes; 1–35 for 150 genomes) peptides was used as database. Peptide identification was performed by DIA-NN under same conditions.

**Supplementary Figure 4.**
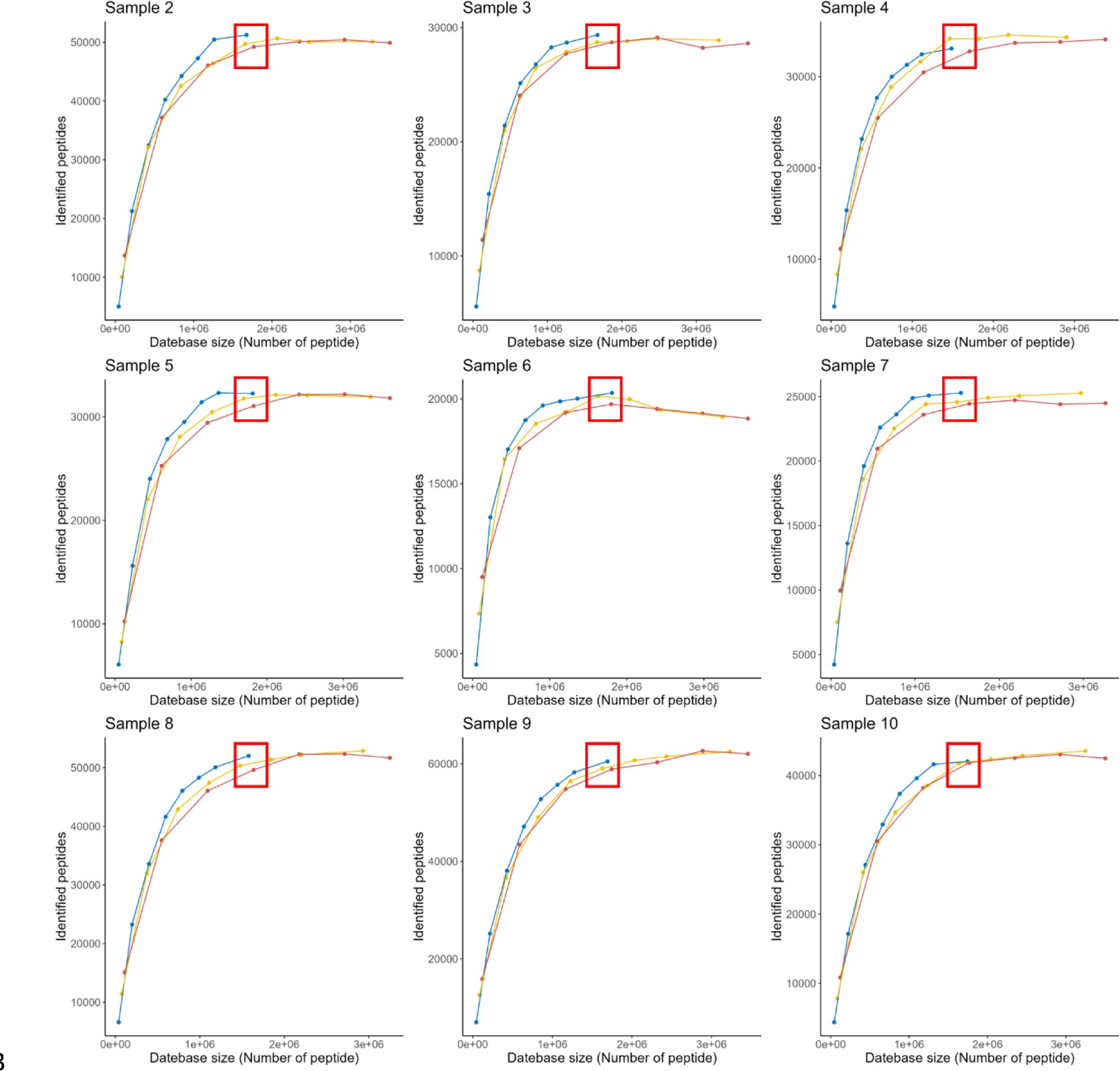
Optimization for genome number and FuncTax score (Sample 2-10). Number of identified peptides against database size. The inflection point has been highlighted with a red box. Peptides from n (50, 100, 150) genomes were ranked by the FuncTax score and top x% (1-40 for 50 and 100 genomes; 1–35 for 150 genomes) peptides was used as database. Peptide identification was performed by DIA-NN under same conditions.

**Supplementary Figure 5.**
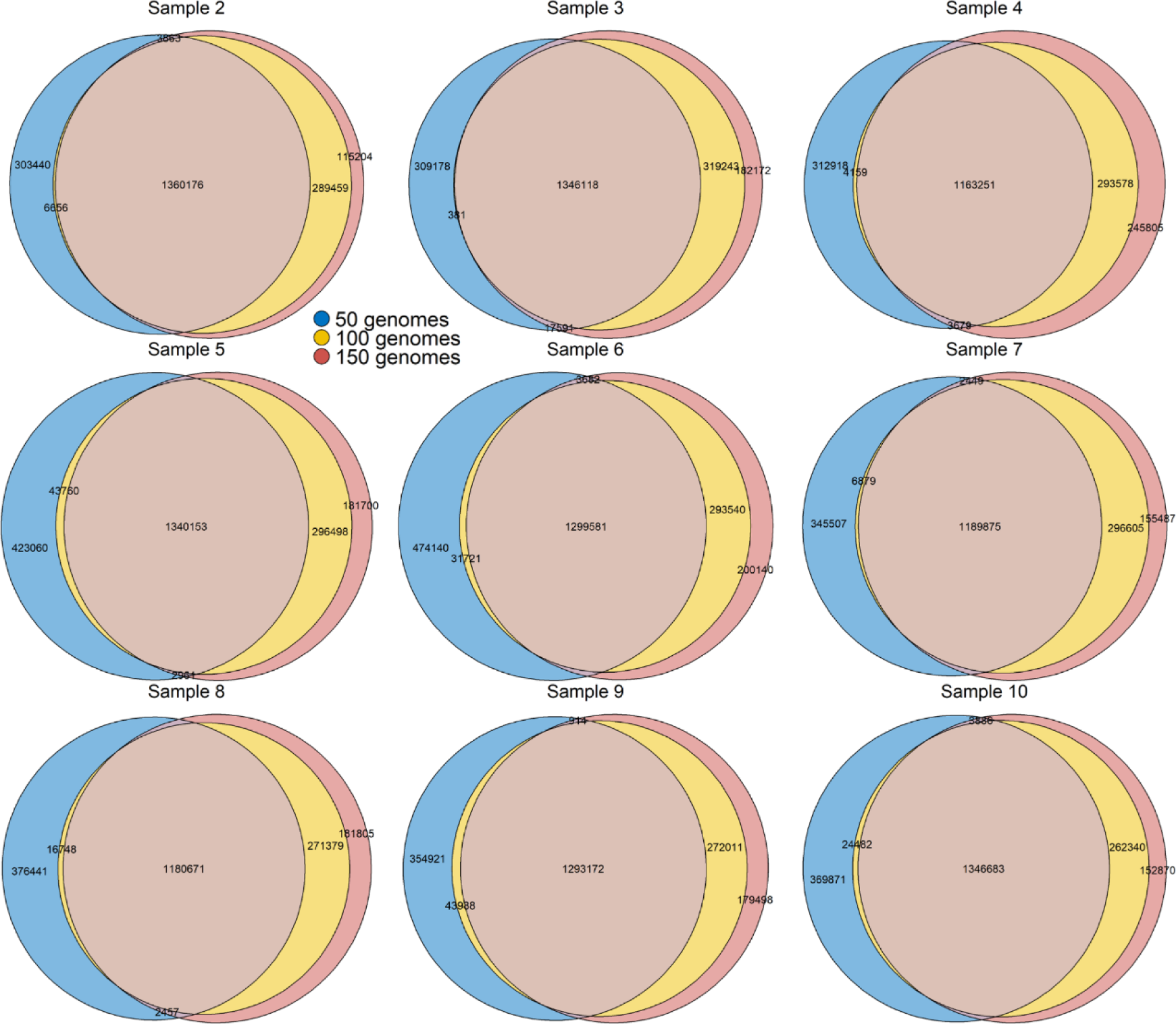
Optimization for genome number and FuncTax score (Sample 2-10). The overlap of reduced peptide database when taking top 40% peptides for 50 genomes, top 20% for 100 genomes and top 15% for 150 genomes as database.

**Supplementary Figure 6.**
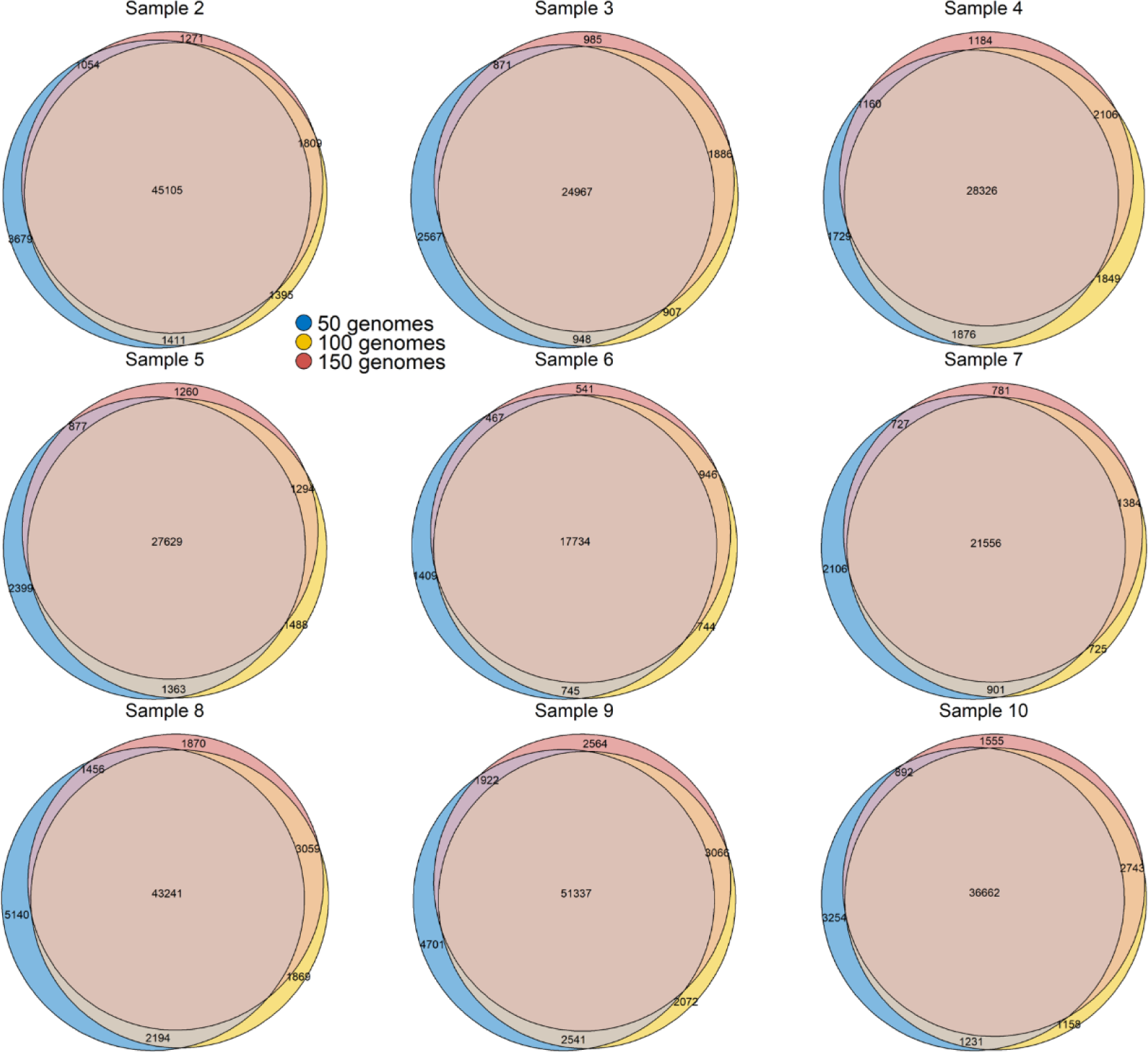
Optimization for genome number and FuncTax score (Sample 2-10). The overlap of identified peptide when taking top 40% peptides for 50 genomes, top 20% for 100 genomes and top 15% for 150 genomes as database. Peptide identification was performed by DIA-NN under same conditions.

**Supplementary Figure 7.**
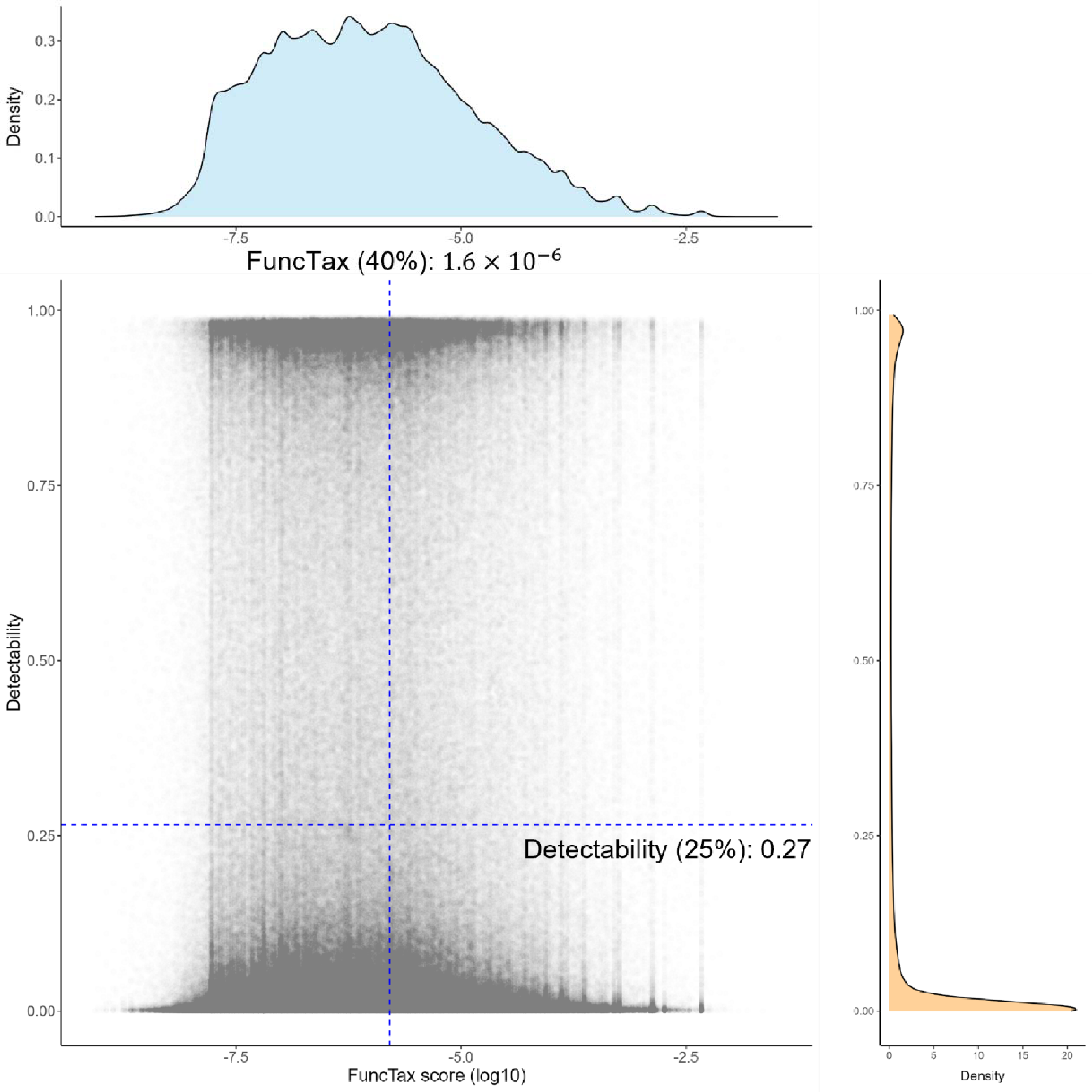
The distribution of detectability score and FuncTax score (Sample 1, top 50 genomes). The dotted blue line shows the cutoff for detectability (top 25%) and FuncTax (top 40%).

**Supplementary Figure 8.**
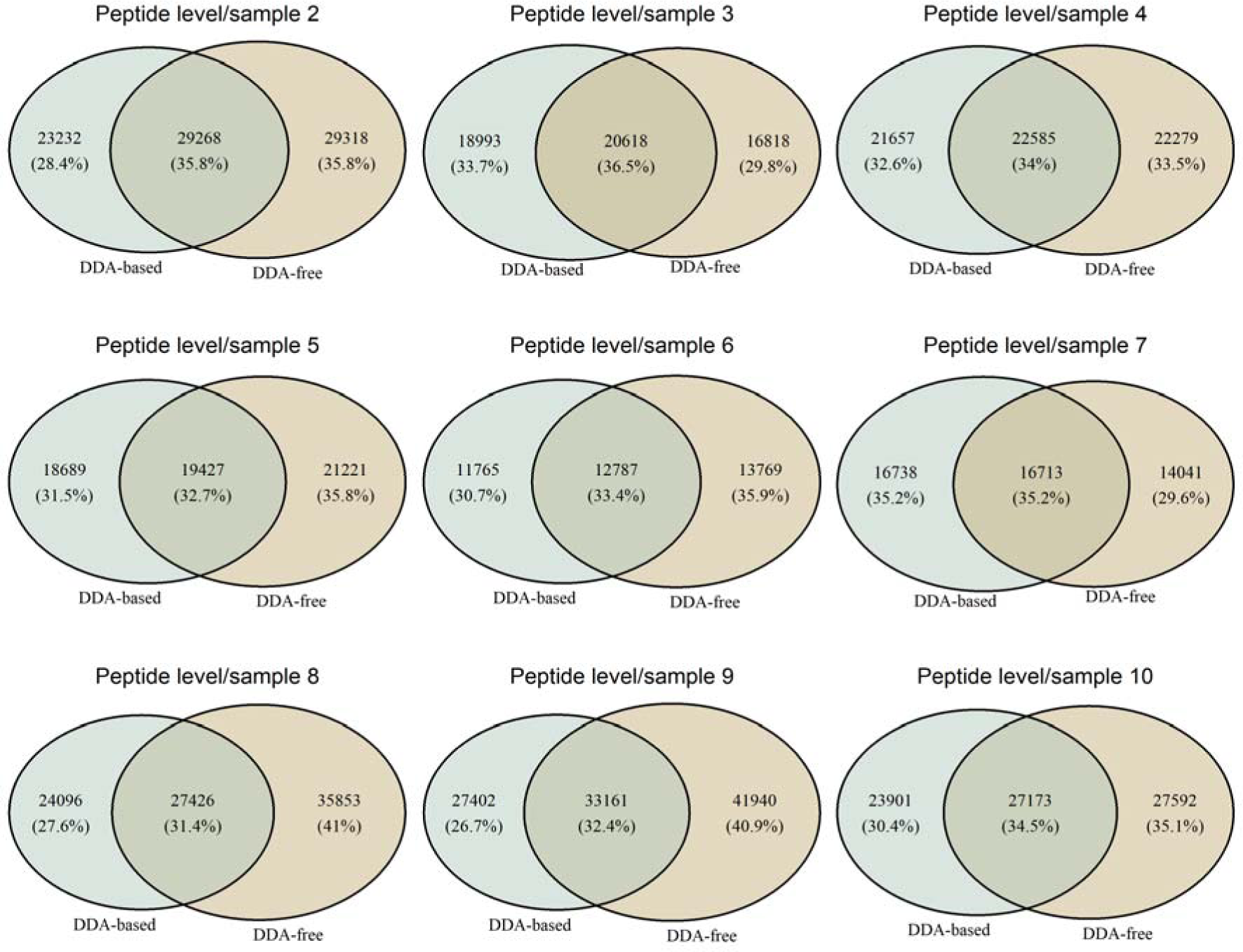
The overlap of peptide identified by each DDA-based method and DDA-free method (Sample 2-10)

**Supplementary Figure 9.**
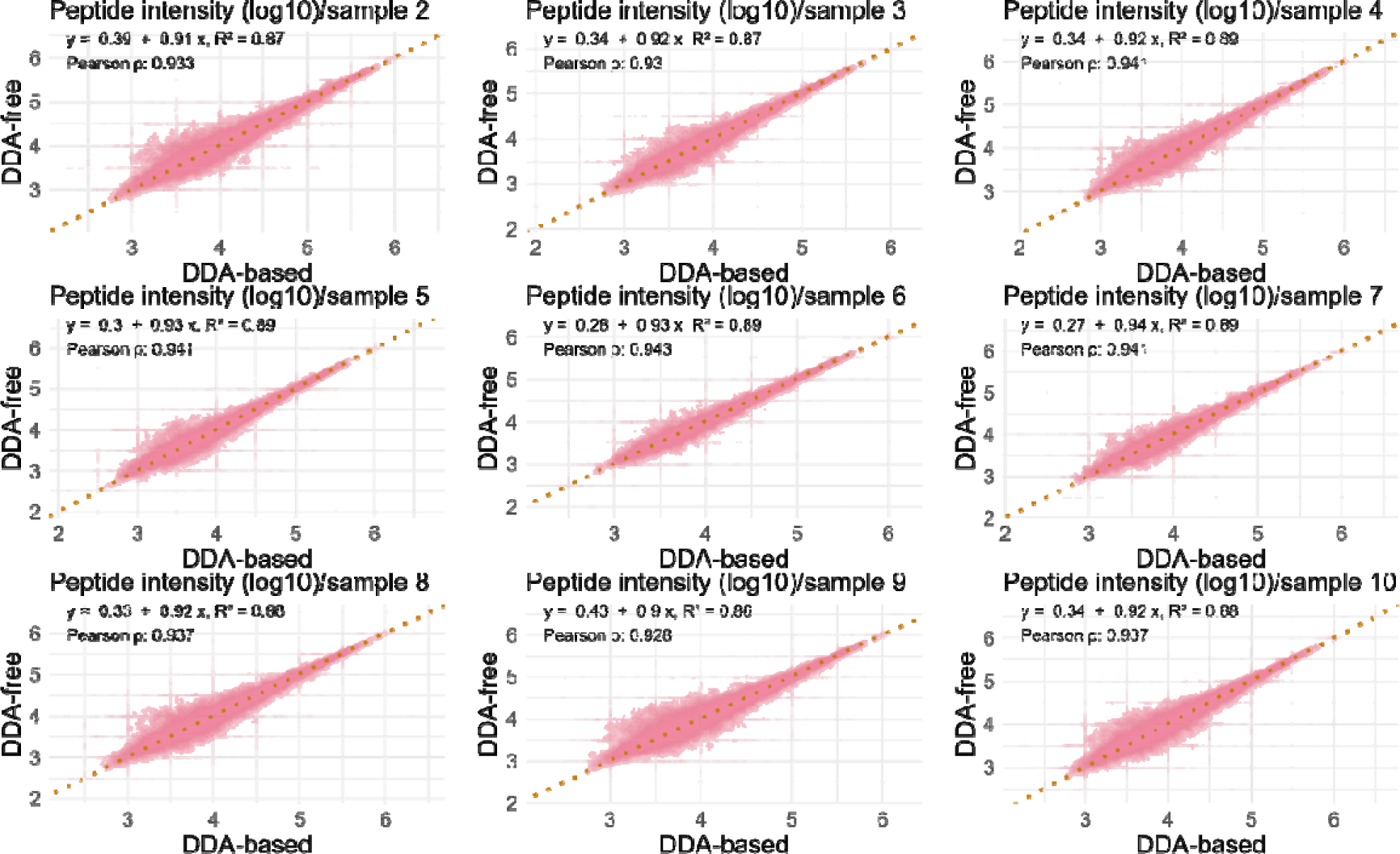
The intensity correlation of the overlapped peptides found by both DDA-based and DDA-free method (Sample 2-10). The dashed line indicates y = x. For DDA-based method, the peptides identified as derived from human proteins are removed.

**Supplementary Figure 10.**
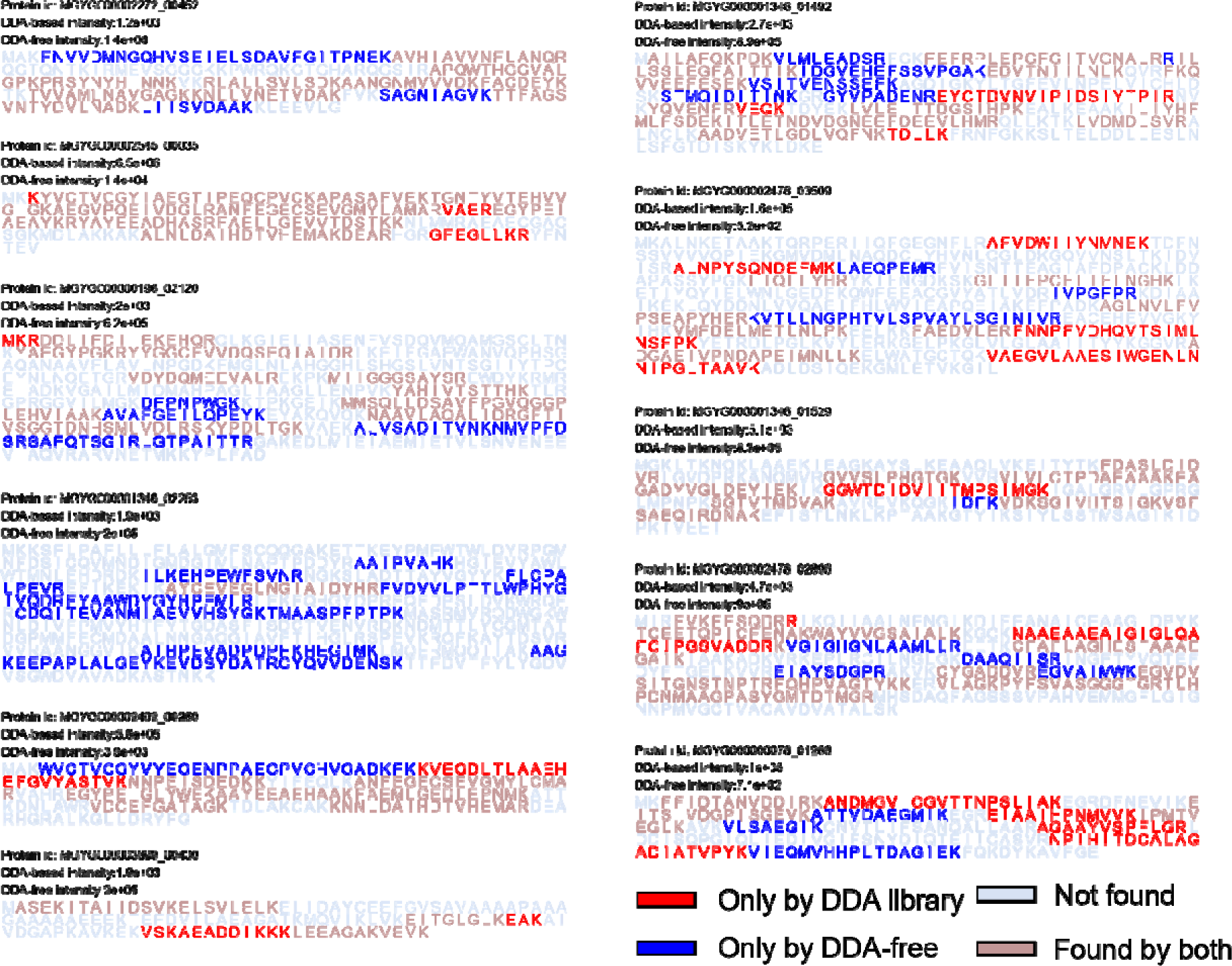
The sequence coverage of representative proteins that exhibit the largest relative difference in intensity DDA-based and DDA-free method.

**Supplementary Figure 11.**
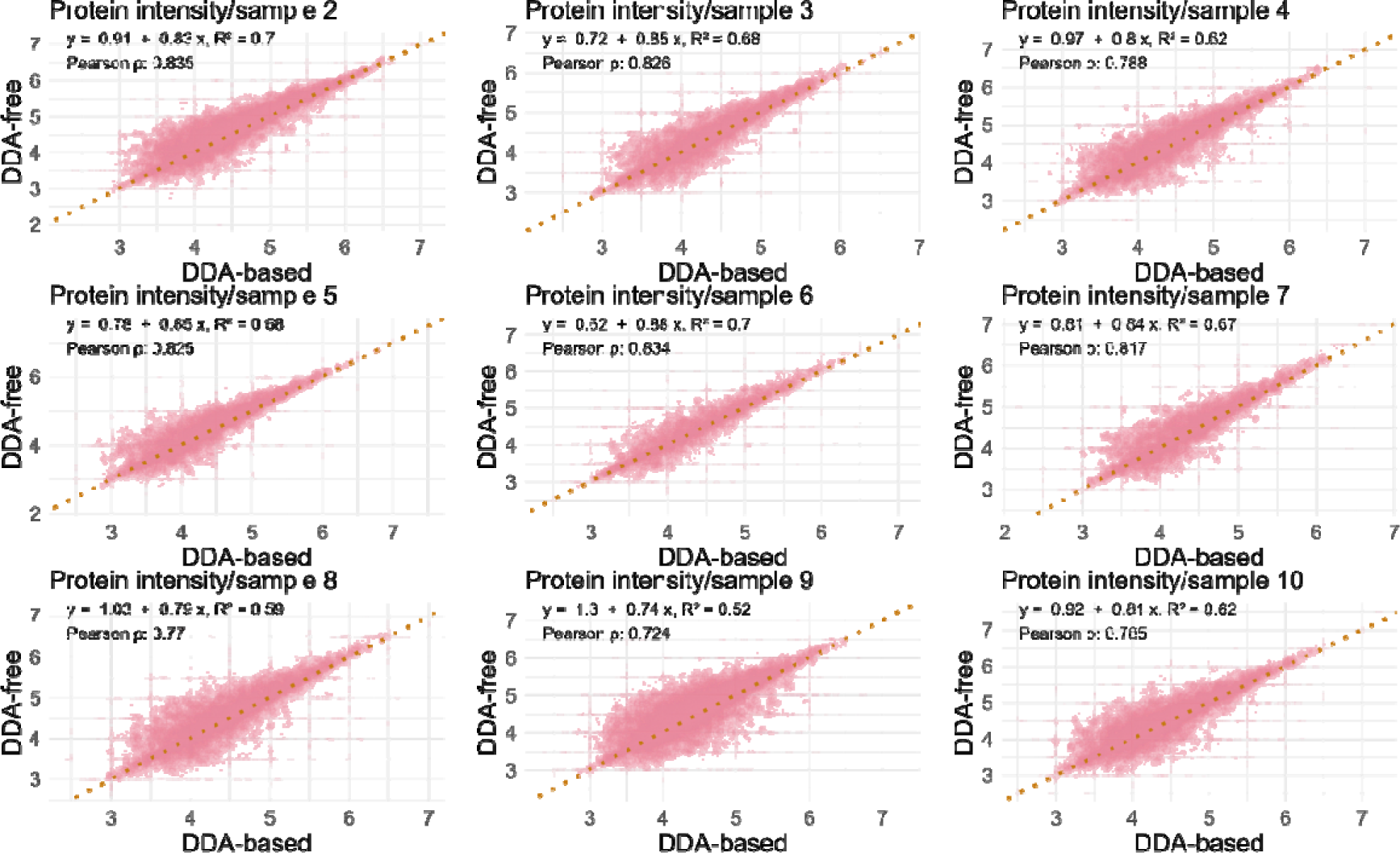
The intensity correlation of the overlapped proteins found by both DDA-based and DDA-free method (Sample 2-10). The dashed line indicates y = x.

**Supplementary Figure 12.**
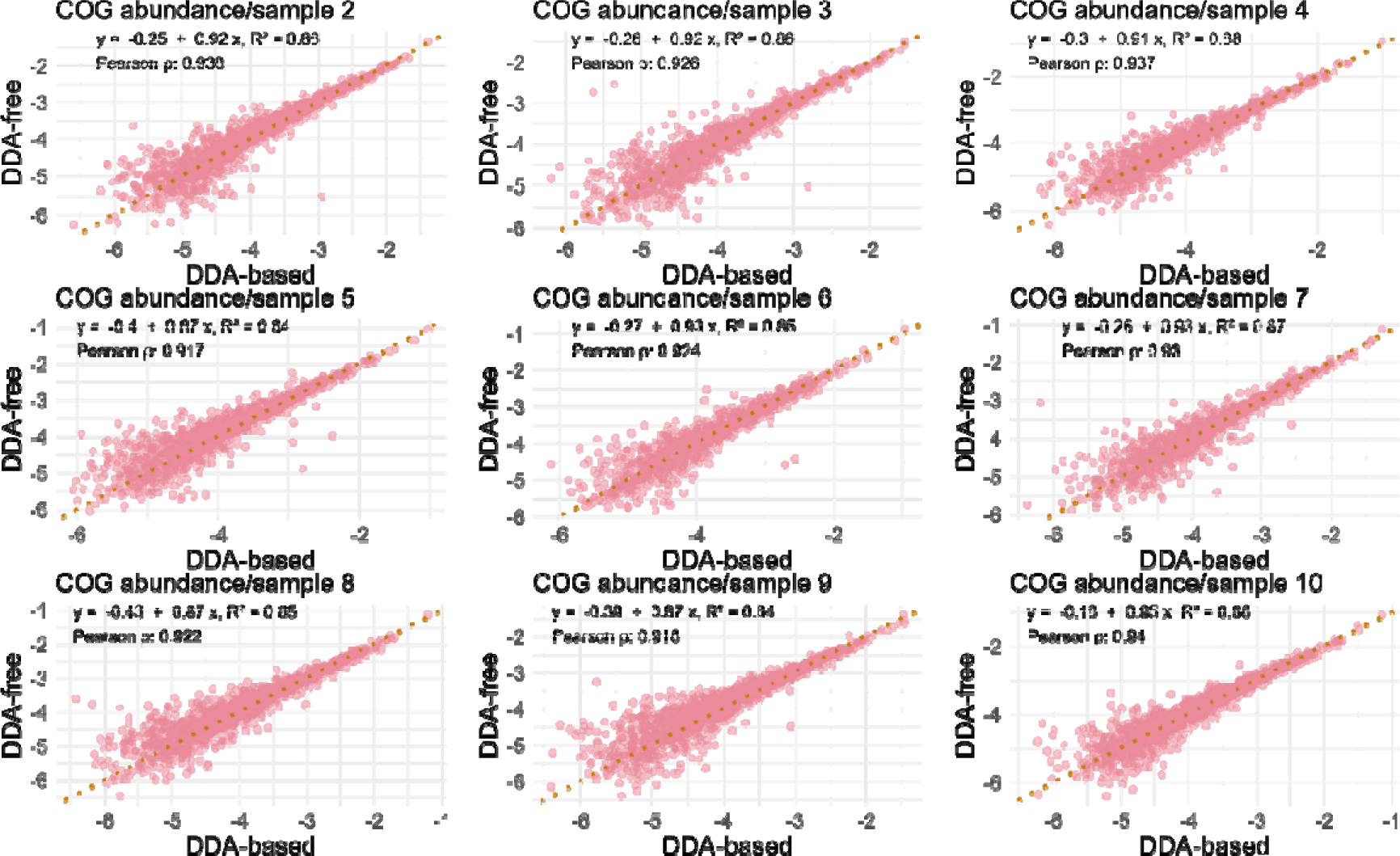
The intensity correlation of the overlapped COG accessions found by both DDA-based and DDA-free method (Sample 2-10). The dashed line indicates y = x.

**Supplementary Figure 13.**
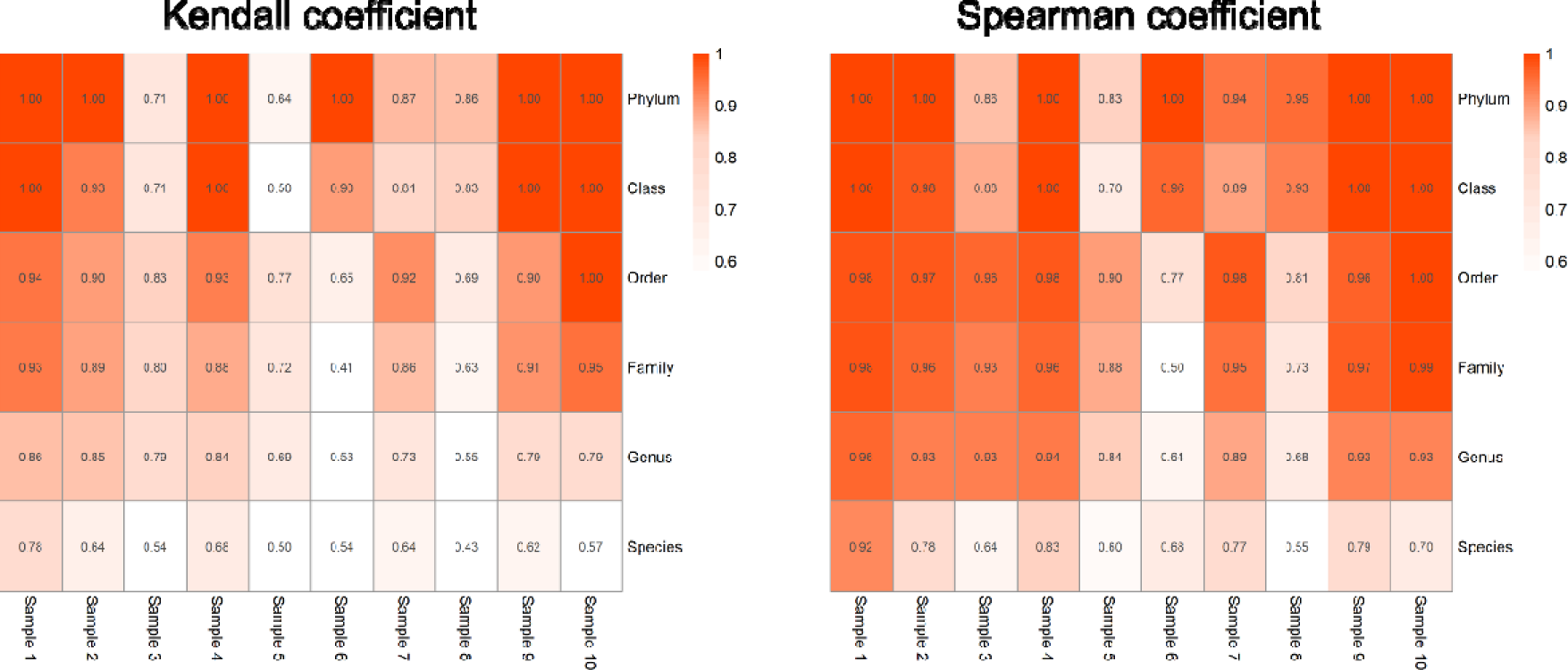
Correlation analysis between DDA-based method and DDA-free method on different taxonomic levels from Species to Phylum. The relative abundance was used for the analysis. Taxonomic categories that were unique to one method were imputed with a value of zero.

**Supplementary Figure 14.**
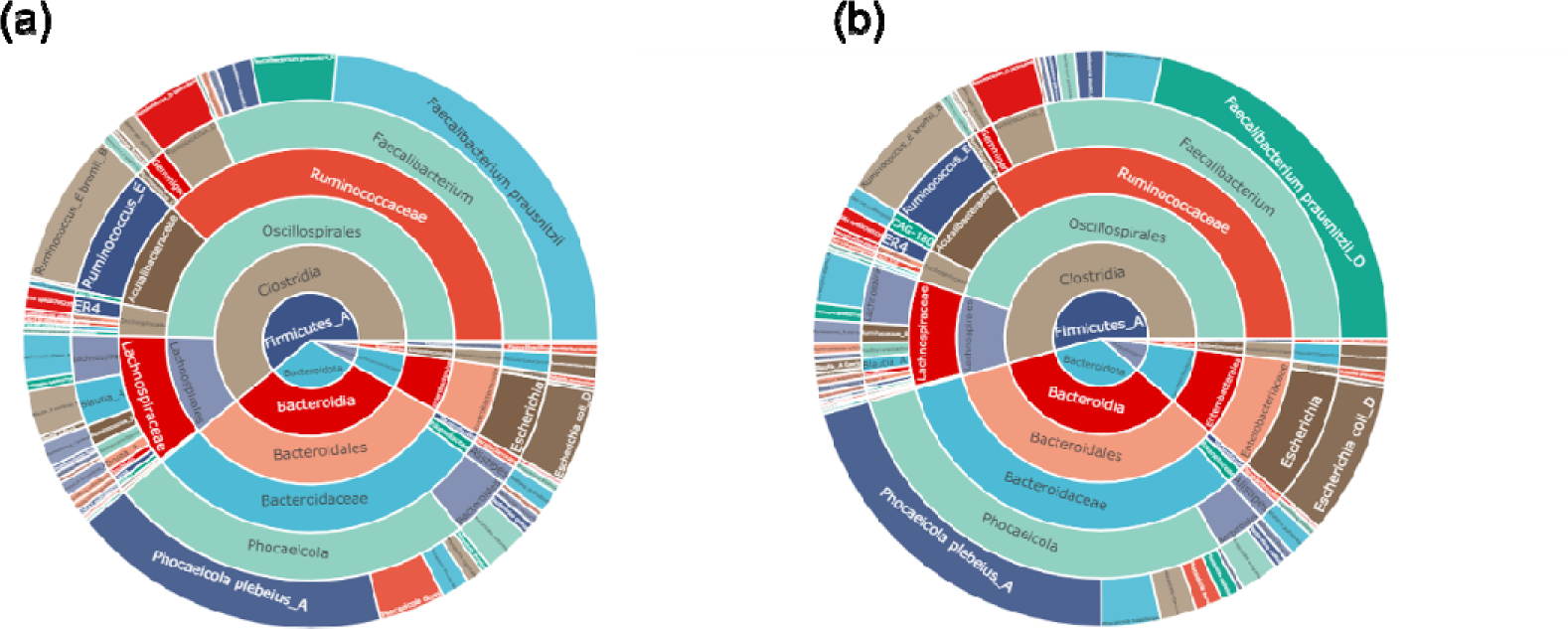
Comparative Taxonomic Composition of the Microbiome at Levels from Phylum to Species. (a) DDA-Based. (b) DDA-Free.

**Supplementary Figure 15.**
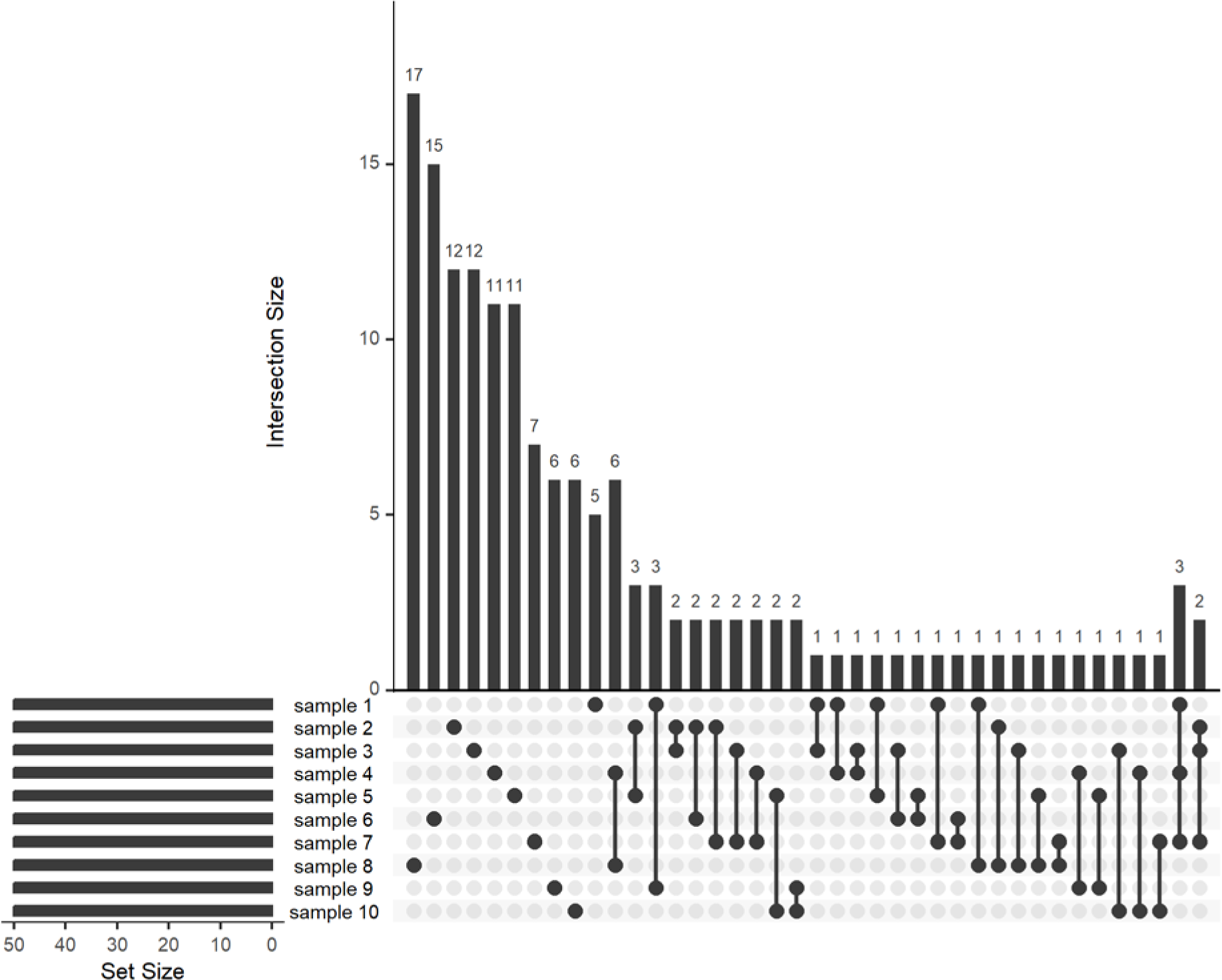
UpSet plot illustrating the overlap in genomes identified by the DDA-free method across ten microbiome samples. The top 40 intersections were plotted.

